# Metabolic Atlas of Early Human Cortex Identifies Regulators of Cell Fate Transitions

**DOI:** 10.1101/2025.03.10.642470

**Authors:** Jessenya Mil, Jose A. Soto, Nedas Matulionis, Abigail Krall, Francesca Day, Linsey Stiles, Katrina P. Montales, Daria J. Azizad, Carlos E. Gonzalez, Patricia R. Nano, Antoni A. Martija, Cesar A. Perez-Ramirez, Claudia V. Nguyen, Ryan L. Kan, Madeline G. Andrews, Heather R. Christofk, Aparna Bhaduri

**Author notes:** These authors contributed equally (Jessenya Mil, Jose A. Soto).

## Abstract

Characterization of cell type emergence during human cortical development, which enables unique human cognition, has focused primarily on anatomical and transcriptional characterizations. Metabolic processes in the human brain that allow for rapid expansion, but contribute to vulnerability to neurodevelopmental disorders, remain largely unexplored. We performed a variety of metabolic assays in primary tissue and stem cell derived cortical organoids and observed dynamic changes in core metabolic functions, including an unexpected increase in glycolysis during late neurogenesis. By depleting glucose levels in cortical organoids, we increased outer radial glia, astrocytes, and inhibitory neurons. We found the pentose phosphate pathway (PPP) was impacted in these experiments and leveraged pharmacological and genetic manipulations to recapitulate these radial glia cell fate changes. These data identify a new role for the PPP in modulating radial glia cell fate specification and generate a resource for future exploration of additional metabolic pathways in human cortical development.

## Introduction

For stem cells to differentiate, proper activation of gene expression and cell signaling must be achieved. Recent advances in studying the developing human cortex have enabled the field to leverage genomic and single-cell methods to begin to understand the cell types that exist during development (Bhaduri et al., 2021; Braun et al., 2023; Nowakowski et al., 2017; Velmeshev et al., 2023). Though recent studies in rodents suggest that metabolism may be an essential driving force through these key transitions (Perez-Ramirez et al., 2024), the metabolism that enables the rapid proliferation and differentiation of the human cortex remains largely unexplored.

### Metabolism Changes with Stem Cell Transitions

Previous studies have elaborated on the role of metabolism as a key regulator for stem cells transitioning into mature and differentiated states. Such studies have described glycolytic metabolism as necessary to maintain a pluripotent state in embryonic stem cells (ESCs), whereas inhibition of glycolysis led to decreased pluripotency signatures and increased differentiation (Gu et al., 2016). However, it has also been observed that high glycolytic activity, through high extracellular glucose levels, may also restrain pluripotent stem cells (PSCs) from reaching a mature state as seen in cardiomyocytes (Nakano et al., 2017). Thus, regulation of metabolism in general, and glycolysis in particular, is important for maintaining homeostasis and stemness. In neurogenesis, neural stem cells (NSCs) have been observed to decrease their use of glycolysis and begin to utilize oxidative phosphorylation (OXPHOS) during neuronal differentiation (Zheng et al., 2016). The reduction of glycolytic activity is imperative during maturation as continuous overexpression of glycolytic genes in NSCs has been shown to decrease neuronal survival (Zheng et al., 2016). In addition, a conditional deletion of mitochondrial genes can impact brain development by reducing the number of NSCs (Khacho et al., 2017). Mechanisms for these actions have primarily linked metabolites with defined roles in modulating epigenetic marks to transcriptional programs resulting in a direct impact on stem cell transitions and cell fate specification (Zhang et al., 2018).

### Cortical Development

Metabolic changes have been well-described as correlated to stem cell transitions and a regulator of these processes in some tissues. However, metabolic changes linked to specific cell types throughout brain development are less fully developed. Here, we interrogate metabolism in the context of human cortical development, which is a dynamic and expansionary process with many iterations of progenitor cell type maturation and diversification. Initially, the human cortex is largely defined by a relatively uniform neuroepithelium that gives rise to radial glia (RG). These RG give rise to neurons through direct and then indirect neurogenesis via transit amplifying intermediate progenitor cells (IPCs). Newborn neurons migrate up the radial glia scaffolding, resulting in the deposition of deep then upper layer neurons before shifting to a gliogenic program that generates astrocytes and oligodendrocyte precursor cells (Cadwell et al., 2019). The major gene programs that correspond to these transitions have been linked to core “developmental periods” (DPs) (Kang et al., 2011) that we use here to contextualize the biological transitions that dictate the peak stages of neurogenesis that we primarily explore. Over several weeks, the developing human cortex expands immensely, suggesting shifts in metabolism across various emerging cell types underpin cortical development and the expansion of the cortex in human evolution (Namba et al., 2021).

### Brain Metabolism

Recent studies have increasingly explored the intersection between metabolism and cortical development, particularly the roles of metabolic gene expression and individual metabolites in rodent models (Namba et al., 2021). For example, mitochondrial dynamics of radial glia were found to be important in determining differentiation into neuronal cells across systems (Iwata et al., 2020; Iwata and Vanderhaeghen, 2021). Additionally, lactate production in radial glia was discovered as a regulator of radial glia division and brain vasculature development in mice (Wu et al., 2023). A metabolic atlas of mid-to-late mouse gestational development found that hyperglycemic pregnancy altered amino acid neurotransmitters in the brain which may suggest a higher incidence of congenital brain defects with increased blood glucose (Perez-Ramirez et al., 2024). Collectively, these studies demonstrate the mitochondrial, cytosolic, and extracellular nutrient availability regulate neuronal differentiation. However, most of our understanding of the relationship between metabolism and early neuronal development comes from research involving rodent models (Namba et al., 2021). There exist important evolutionary differences between mouse and human cortical development such as the existence and behavior of cell types like outer radial glia (oRG) and IPCs with enrichment in primates and subtype diversification (Betizeau et al., 2013; Lui et al., 2011). These evolutionary differences also extend into metabolism as studies have manipulated human-specific metabolic genes including ARGHAP11B, a mitochondrial gene, and TKTL1, a human-specific version of a pentose phosphate pathway gene in rodents. These experiments have shown that ARGHAP11B and TKTL1 can drive IPC and oRG expansion in rodents, mimicking behavior of human brain progenitors (Namba et al., 2020; Pinson et al., 2022). Moreover, mitochondrial metabolism is important in regulating the faster pace of neuronal development in mice compared to humans (Iwata et al., 2023). Despite the advances in understanding neurometabolism, the metabolic processes controlling human cortical development, particularly during early stages when sample acquisition is limited, remain poorly understood.

### Cortical Organoids and Role of Metabolism in Cell Fate Specification

Cortical organoids have become a valuable tool in studying human cortical development. As pluripotent stem cell-derived models of the developing human brain, cortical organoids generally recapitulate the major cell types and broad developmental transitions observed in primary tissue (Camp et al., 2015; Pollen et al., 2019; Velasco et al., 2019). As a tool to interrogate the role of metabolism in modulating human cortical development, they are an emerging resource. Some studies have indicated an altered glycolytic gene expression program as being a major difference between cortical organoids and primary tissue (He et al., 2024) that impacts appropriate cell fate specification (Bhaduri et al., 2020), while others have contended this is simply an artifact of the system (Vertesy et al., 2022) that does not impair fate acquisition (Uzquiano et al., 2022). However, across studies, no comprehensive characterization of metabolites has yet been performed, nor have these metabolomic abundances been benchmarked to the primary context. Moreover, cortical organoids are an alluring model in which to study the biological underpinnings of the relationship between metabolism and fate specification as they are scalable, easily tractable with environmental, pharmacological, or genetic modulations, and can be readily characterized across developmental time.

To understand how metabolism changes during and drives human cortical development, we have generated a metabolic atlas of development in both primary tissue and in cortical organoids. Using this atlas, we identify highly dynamic known and novel changes in key metabolic pathways. Based on these observations, we were led to explore how glycolysis, through the pentose phosphate pathway (PPP), impacts cell fate specification. Using a combination of this atlas and functional experiments in cortical organoids, we find that PPP activity peaks during developmental period 4 (DP4), inhibits radial glia maturation and modulates radial glia cell fate specification. This highlights how metabolism regulates cell fate in the context of the human cortex and demonstrates how our atlas and cortical organoids can be used to uncover these foundational features of human cortical cell fate diversification.

## Results

### Leveraging Plated Human Primary Tissue to Generate a Metabolic Atlas

To generate an atlas of human cortical development, we dissociated and plated developmental post-mortem human primary cortical tissue (HPCT) onto coated plates (Methods) in order to perform immunofluorescence (IF), Seahorse, and liquid chromatography mass spectrometry (LCMS)-based metabolomics and metabolic tracing measurements (Fig 1A). This approach to studying HPCT is widely used and has been observed to recapitulate key cell types and biological processes found in primary samples (Andrews et al., 2020; Ostrem et al., 2014). For these analyses, we leveraged 19 primary tissue samples spanning 5 developmental periods aligned to previously described developmentally important stages (Kang et al., 2011) (SFig 1A). We decided to leverage a two-dimensional primary culture approach to allow for both Seahorse Assay and the ^13^C_6_-glucose and ^13^C_5_-glutamine tracing measurements that would be inaccessible with immediate analysis; these approaches have been used previously to identify interesting processes in understanding the cortical development and changes in the developing mouse brain in a diabetic model (Dong et al., 2022; Perez-Ramirez et al., 2024). Additionally, this plating strategy enriches for progenitors and their locally produced daughter neurons and glia as more mature cells are unlikely to survive the plating, allowing for an emphasis on the study of cell fate acquisition metabolic programs.

**Figure 1.**
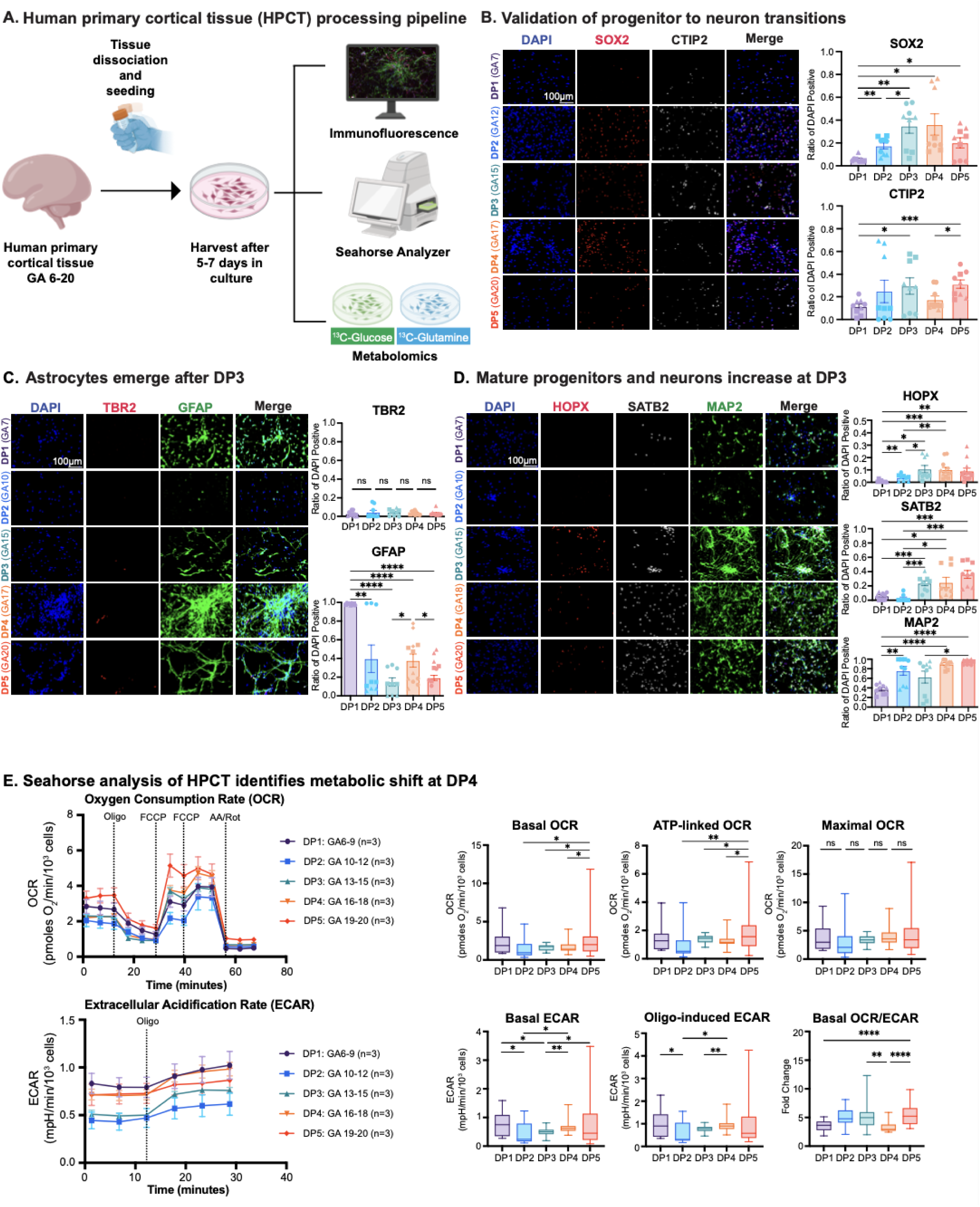
Human Primary Cortical Tissue Processing Pipeline Enables Metabolic Measurements Across Developmental Periods. A) Schematic of Human Primary Cortical Tissue (HPCT) processing pipeline. Primary tissue from gestational age (GA) 6 - 20 are obtained from clinical sources with anonymized identity and appropriate consent. To create a metabolic atlas, including functional measures of metabolism, we dissociated and plated the primary cells. After 5 - 7 days in culture, immunofluorescence (IF) was used to evaluate cell-type composition, Seahorse Analysis evaluated functional glycolysis and oxidative processes, and metabolomics was performed for both total abundance and ^13^C_6_-glucose and ^13^C_5_-glutamine tracing. A total of 28 samples were characterized using this pipeline, with 10 having matched data across all modalities and 27 samples having at least 2 modalities measured. B) To validate that cell-type composition mirrored expected patterns in HPCT, we performed IF. A representative image for each of the 5 developmental periods (DP) is shown here. Staining for SOX2 (red) was used as a proxy for radial glia progenitor identity, whereas staining for CTIP2 (white) labeled emerging deep layer neuron populations. DAPI, shown in blue, marked nuclei. Scale bar is 100 μM and reflects all images. Quantification of the percent of DAPI positive cells marked by each protein is presented in bar charts to the right, where 3 distinct biological replicates were analyzed for each developmental period (DP), and 3 technical replicates were used within each biological replicate; shapes on the bar chart correspond to individual biological replicates within a DP. DP1 corresponds to GA 6 - 9, DP2 corresponds to GA 10 - 12, DP3 corresponds to GA 13 - 15, DP4 corresponds to GA 16 - 18, and DP 5 corresponds to GA 19 - 20. Significance was calculated using the Brown-Forsythe and Welch ANOVA. * indicates p-value < 0.05, ** indicates p-value < 0.01, *** indicates p-value < 0.001. C) Staining and quantification for TBR2 (red, gene *EOMES*) which marks intermediate progenitor cells, and GFAP (green) which marks radial glia early and astrocytes later, was performed as in (B). The distinction between GFAP (green) marking radial glia versus astrocytes can be visualized via cell morphology. DAPI, shown in blue, marked nuclei. Scale bar is 100 μM and reflects all images. A representative image for each of the 5 developmental periods is shown here, with n = 3 biological replicates for DPs 1 - 3 and 5, and n = 4 for DP4. 3 technical replicates were used for each biological replicate. Significance was calculated using the Brown-Forsythe and Welch ANOVA. Non-significant trends marked by ns. * indicates p-value < 0.05, ** indicates p-value < 0.01, **** indicates p-value < 0.0001. D) Staining and quantification for HOPX (red) which marks outer radial glia, SATB2 (white) which marks upper layer neurons, and MAP2 (green) which is a pan-neuronal marker was performed as in (B). DAPI, shown in blue, marked nuclei. Scale bar is 100 μM and reflects all images. A representative image for each of the 5 developmental periods is shown here, with n = 3 biological replicates for DPs 1 - 3 and 5, and n = 4 for DP4 and DP5 for HOPX staining only. 3 technical replicates were used for each biological replicate. Significance was calculated using the Brown-Forsythe and Welch ANOVA. * indicates p-value < 0.05, ** indicates p-value < 0.01, *** indicates p-value < 0.001, **** indicates p-value < 0.0001. E) Seahorse analysis was performed to measure oxygen consumption rate (OCR, top line graph) and extracellular acidification rate (ECAR, bottom line graph). Features of these graphs are presented in box and whisker plots on the right. Basal OCR measures the initial OCR of each sample. During the Seahorse analysis, small molecule inhibitors are delivered at uniform timepoints during the experiment in order to elicit specific responses and quantitative measures therein. For OCR, oligomycin (oligo) inhibits ATP-synthase, enabling measurement of ATP-linked OCR. Maximal OCR is calculated after the delivery of carbonyl cyanide-p-trifluoromethoxyphenylhydrazone (FCCP) which works as an uncoupler. Basal ECAR is a proxy for glycolysis and measures the baseline glycolytic rate, while after oligo Oligo-induced ECAR is measured. The Basal OCR/ECAR is a calculated feature. Both OCR and ECAR are normalized to the number of cells in the plate, which is determined based upon Hoescht staining and plate-based, high-content imaging. 3 biological replicates of each DP were analyzed, with 6 - 12 technical replicates for each sample. A significant decrease in the basal OCR/ECAR ratio was observed at DP4. Significance was calculated with an ANOVA and a Tukey’s multiple comparisons test. ** indicates p-value < 0.01, **** indicates p-value < 0.0001.

We first validated that each sample used in our metabolic atlas generation pipeline was cortical by performing quantitative real-time PCR (qRT-PCR) for *FOXG1* and *BCL11B* to verify cortical identity; all samples included in this study were confirmed to be cortical (SFig 1B) (Leone et al., 2008; Rubenstein, 2000). To verify that cell type composition in these two-dimensional primary cultures preserved the major cell types and proportions observed *in vivo*, we performed extensive immunofluorescence staining and quantification across developmental periods. Across the analysis, the younger samples had more highly expressed progenitor and early neuronal features (Fig 1B), while older DPs expressed markers of emerging astrocytes (Fig 1C), more mature progenitors such as outer radial glia (oRG, (Pollen et al., 2015)) and upper layer neurons (Fig 1D) as expected.

### Indirect Measures of Glycolysis and Oxidative Phosphorylation Identify Increased Reliance on Glycolysis at Developmental Period 4

Having validated our system’s ability to recapitulate expected cell types and developmental transitions, we performed Seahorse analysis to assess metabolic shifts across DPs. Seahorse is used as an indirect measure of glycolysis by measuring extracellular acidification (ECAR) and oxidative phosphorylation (OXPHOS) through measurement of the oxygen consumption rate (OCR) (Divakaruni et al., 2014). We were encouraged that within a DP, replicates were very consistent given the variability that is often associated with HPCT procurement and processing (SFig 1C). From our results we saw that basal OCR and ATP-linked OCR increased significantly across developmental periods (Fig 1E) which has been described as a metabolic transition that occurs when differentiating from neural stem cells to neurons (Zheng et al., 2016), likely due increasing energy demands. We expected the ECAR, which indirectly measures glycolysis, would decrease over time. Generally, this downward trend was observed (Fig 1E), but our data shows that glycolytic activity is more dynamic over cortical development than we previously thought.

Overall, the observed OCR and ECAR trends fit our prediction of a progenitor-to-mature transition across developmental stages. However, we see an increased dependence on ECAR during DP4 which corresponds to when oRG cells are at their maximal abundance and when the onset of gliogenesis begins and upper layer neurogenesis is ongoing (Cadwell et al., 2019). This was specifically emphasized by the significant decrease in the OCR/ECAR ratio at DP4 (Fig 1E). However, many of these developmental transitions have already begun by DP3 (Cadwell et al., 2019), suggesting unique metabolic features of DP4 that cannot be fully explained by cell-type composition. These intriguing dynamics motivated us to further explore HPCT metabolism by performing metabolomic analysis.

### Metabolomics of HPCT Shows Dynamic Metabolite Shifts Across Developmental Time

To more directly measure metabolites and metabolic pathway activities in our HPCT cells, we performed LCMS-based metabolomics and metabolic tracer analysis. HPCT cells were dissociated and plated for 5 days before introduction of ^13^C_6_-glucose and ^13^C_5_-glutamine tracer for 24 hours. Metabolites were then harvested from the HPCT cells, matching the parallel IF and Seahorse harvest timepoints. Extracted metabolites were analyzed with LCMS and analyzed against a library of known metabolites as has been recently described (Perez-Ramirez et al., 2024). For this dataset, we mapped to 126 metabolites (Fig 2A, STables 1 and 2) from the validated library and also generated data related to isotopologue composition from the traced metabolites (STable 3).

**Figure 2:**
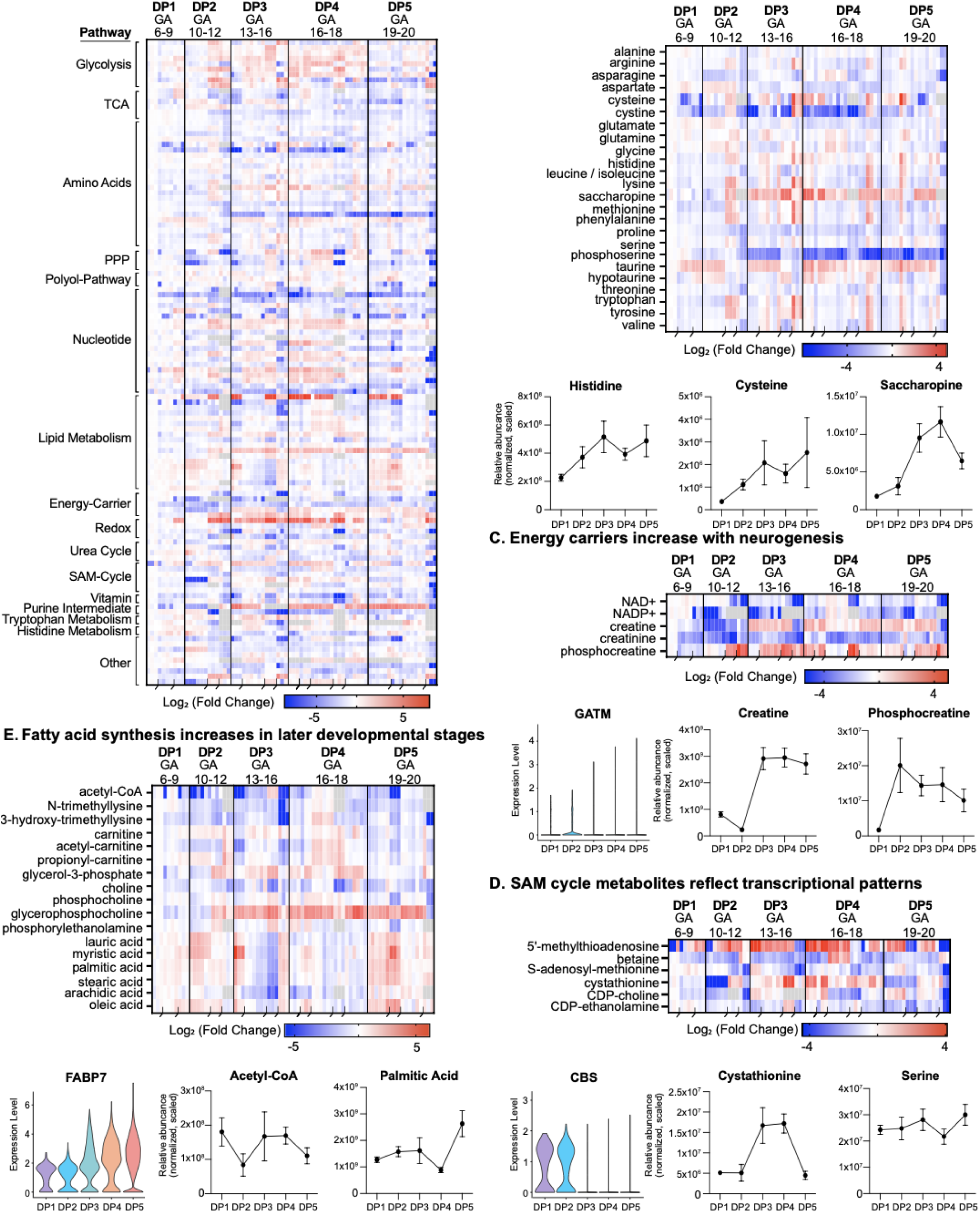
Metabolomics Identifies Dynamic Pathways Across Human Cortical Development. A) Heatmap of 127 targeted metabolite abundances throughout cortical development organized by pathway. Metabolomics was performed on plated HPCT using liquid chromatography mass spectrometry (LCMS) to measure baseline metabolite levels. Pooled metabolite values were normalized by DNA concentration and trifluoromethanesulfonate (TFMS). Each metabolite measurement was scaled to an arbitrary number for data representation. Metabolite levels represent a log_2_ fold change to the youngest sample collected (GA6). 3 - 5 biological replicates of each DP were analyzed, with 3 - 6 technical replicates for each sample. Blue or red values indicate a negative or positive log fold change to GA6, respectively. Gray values represent an absence of data collected for a metabolite from the metabolomics analysis. Black indents at bottom of heatmap separate biological replicates within each DP. B) Heatmap of amino acids levels across cortical development and metabolite abundance graphs of histidine, cysteine and saccharopine. Metabolite levels in the heatmap represent a log_2_ fold change to GA6. Three to five biological replicates of each DP were analyzed, with three to six technical replicates for each sample. Combined metabolite abundance values for histidine, cysteine and saccharopine were calculated by averaging all technical replicates across biological samples within each DP. Y-axis measures abundance (measured and scaled) across each DP. Error bars represent standard error of mean (SEM). C) Heatmap of energy carrier levels across cortical development, transcriptional expression of glycine amidinotransferase (*GATM*) and metabolite abundance graphs of creatine and phosphocreatine. Metabolite levels in the heatmap represent a log_2_ fold change to GA6. Three to five biological replicates of each DP were analyzed, with three to six technical replicates for each sample. Violin plot for *GATM* was obtained from single-cell RNA sequencing (scRNA-seq) data from (Nano et al., 2023). Y-axis measures normalized and scaled mRNA expression levels across DPs. Combined metabolite abundance values for creatine and phosphocreatine were calculated by averaging all technical replicates across biological samples within each DP. Y-axis measures abundance (measured and scaled) across each DP. Error bars represent SEM. D) Heatmap of SAM cycle metabolite levels throughout cortical development, transcriptional expression of cystathionine beta synthase (*CBS*) and metabolite abundance graphs of cystathionine and serine. Metabolite levels in the heatmap represent a log_2_ fold change to GA6. Three to five biological replicates of each DP were analyzed, with three to six technical replicates for each sample. Violin plot for *CBS* was obtained from scRNA-seq data from Nano et. al 2023 (Nano et al., 2023). Y-axis measures normalized and scaled mRNA expression levels across DPs. Combined metabolite abundance values for cystathionine and serine were calculated by averaging all technical replicates across biological samples within each DP. Y-axis measures abundance (measured and scaled) across each DP. Error bars represent SEM. E) Heatmap of lipid metabolism related metabolites during cortical development, transcriptional expression of fatty acid binding protein 7 (*FABP7*) and metabolite abundance graphs of acetyl-CoA and palmitic acid. Metabolite levels in the heatmap represent a log_2_ fold change to GA6. Three to five biological replicates of each DP were analyzed, with three to six technical replicates for each sample. Violin plot for *FABP7* was obtained from scRNA-seq data from (Nano et al., 2023). Y-axis measures normalized and scaled mRNA expression levels across DPs. Combined metabolite abundance values for acetyl-CoA and palmitic acid were calculated by averaging all technical replicates across biological samples within each DP. Y-axis measures abundance (measured and scaled) across each DP. Error bars represent SEM.

In the pooled analysis by which metabolite abundance is measured without regard to labeled carbons, we identified metabolites across 15 pathways that exhibited dynamic changes across DPs by comparing the fold change of metabolites across time to the earliest time point in our analysis (DP1) (SFig 2A). As with the Seahorse analysis, the replicates between HPCT in the same DPs were consistent (Fig 2A). To explore our metabolic atlas, we looked at specific pathways with interesting changes in metabolite abundance across human cortical development (SFig 2A) and also linked these pathways to known examples that have been previously studied. When compared to gene expression, we noted in some cases the metabolite abundances corresponded to gene expression changes including in previously described metabolically linked gene expression programs (Dong et al., 2022), but in many cases the gene expression and metabolites diverged (SFig 2B). For example, fatty acids increase in the metabolomics data, but the genes decrease during the same DPs (SFig 2B, Fig 2), suggesting that gene expression is not a comprehensive tool to understand metabolomic dynamics developmentally and highlighting the utility of knowing how metabolites change over time.

### Amino Acid Metabolism

We observed that many metabolites related to amino acid metabolism changed across time, with increases or dips in abundance specific to an individual DP (Fig 2B). For example, histidine, cysteine, and saccharopine all largely increased through development in our data. Histidine is a precursor for the histamine neurotransmitter utilized in both brain and spinal cord (He and Wu, 2020), and its increase corresponds to the maturation of neuronal populations in the cortex. Similarly, cysteine is a precursor for hydrogen sulfide, a neurotransmitter that modulates synaptic plasticity and neuronal activity (He and Wu, 2020). Additionally, it has been observed that the degradation of saccharopine is imperative for neuronal development, as an abnormal accumulation of saccharopine leads to impaired dendritic arborization of neurons (Guo et al., 2022). In our data, we observe an increase of saccharopine up to DP4 after which it decreases, consistent with when there is an increase in neurons during the timepoints sampled.

### Energy Carrier Metabolism

We also explored energy carrier metabolism and found that creatine abundance increased after DP2 while phosphocreatine decreased (Fig 2C), consistent with the biochemical conversion of phosphocreatine into creatine which is used when sudden shifts in oxygen or glucose availability occur (Chen et al., 2023). To relate the metabolic dynamics of our dataset to changes in transcriptional expression of the metabolic genes associated with our targeted metabolites, we leveraged the data from a single-cell RNA sequencing meta-atlas of HPCT (Nano et al., 2023). We looked at the expression of GATM, a gene responsible for the precursor of creatine, and saw that its expression spiked at DP2 matching the increase in creatine abundance.

### SAM Cycle Metabolism

Another pathway of interest that showed dynamic changes is the S-adenosyl-methionine (SAM) Cycle. In methionine metabolism, SAM is key metabolite that contributes to a variety of key biological processes including epigenetic regulation, nucleotide biosynthesis, and membrane lipid homeostasis (Lauinger and Kaiser, 2021). From the LCMS based metabolomics, cystathionine, a precursor to the SAM cycle, increased after DP2 and follows the transcriptional peak of cystathionine-β-synthase (CBS), the gene responsible for its production (Fig 2D). Previously, CBS has been described to be expressed in neuroepithelial cells and radial glia during developmental stages (Enokido et al., 2005). CBS uses serine to convert homocysteine into cystathionine, a precursor for cysteine which we saw also increased over time from our previous investigation of amino acid metabolism (Fig 2B and 2D).

### Lipid Metabolism

Consistent with previous findings, fatty acid synthesis increased during later stages of cortical development, coinciding with exit from the stem cell stage (Maffezzini et al., 2020) (Fig 2E). This also aligns with our analysis of *FABP7* gene expression, a fatty acid binding protein (De Rosa et al., 2012) and marker of oRG cells (Pollen et al., 2015) which also increase in abundance over time. *FABP7* is important for neural progenitor cell differentiation and migration (Maffezzini et al., 2020). We observed that acetyl-CoA, a common precursor of fatty acid synthesis, was relatively stable across DPs but that individual fatty acids increased in prevalence, including palmitic acid which is highly abundant in the brain and participates in numerous signaling pathways (Smith and Bazinet, 2024). Each of these examples, ranging from amino acid metabolism to lipid metabolism, largely show expected changes of individual metabolites along with additional context for how the broader pathway changes in human cortical development. These analyses provide confidence that the rest of the atlas can be reliably used for the discovery of novel biology.

### Atlas of Cortical Organoid Metabolism

Dynamic transitions in primary tissue allude to potential roles for metabolic regulation of key biological processes and cell fate transitions. However, experiments in primary samples are challenging and time restricted, and cortical organoids are now being widely used to study human cortical developmental questions (Jourdon et al., 2023; Paulsen et al., 2022). Previous studies using transcriptomic approaches have indicated that the gene expression of metabolic programs in cortical organoids are divergent from primary tissue (Bhaduri et al., 2020; He et al., 2024), though follow up studies suggest this transcriptional difference may not be impactful towards cell fate specification (Uzquiano et al., 2022; Vertesy et al., 2022). To explore which metabolic pathways are preserved or distinct in organoids across developmental time compared to HPCT, we generated a parallel metabolic atlas of cortical organoids.

As in primary tissue, we characterized key features of organoid metabolism across cortical organoids generated from three lines (hESC and iPSC, male and female) at Weeks 5, 8, 10, 12, and 15. Some features of glycolysis and OXPHOS were preserved, including a general increase in Basal OCR, though an important distinction was that ECAR increased over time in organoids (SFig 3A), while it had a specific spike at DP4 in HPCT (Fig 1E). We also performed LCMS-based metabolomics analysis, including incorporating a ^13^C_6_-glucose and ^13^C_5_-glutamine tracer in our organoid culture for 24 hours prior to harvest. We used the same metabolite extraction methods and analytical pipelines yielding data from 123 metabolites from the library (Fig 3A, STable 4-6). We used immunofluorescence to validate cortex identity in organoids and additionally characterize proper progenitor to neuron transitions to match that of primary tissue (Fig 3B, SFig 3B). To support the pairing of cortical organoid age to HPCT, we performed a methylation array analysis and subsequent methyl age analysis pipelines which use beta values to determine and match the percentage of methylated sites from a DNA sample to a reference (Steg et al., 2021).

**Figure 3:**
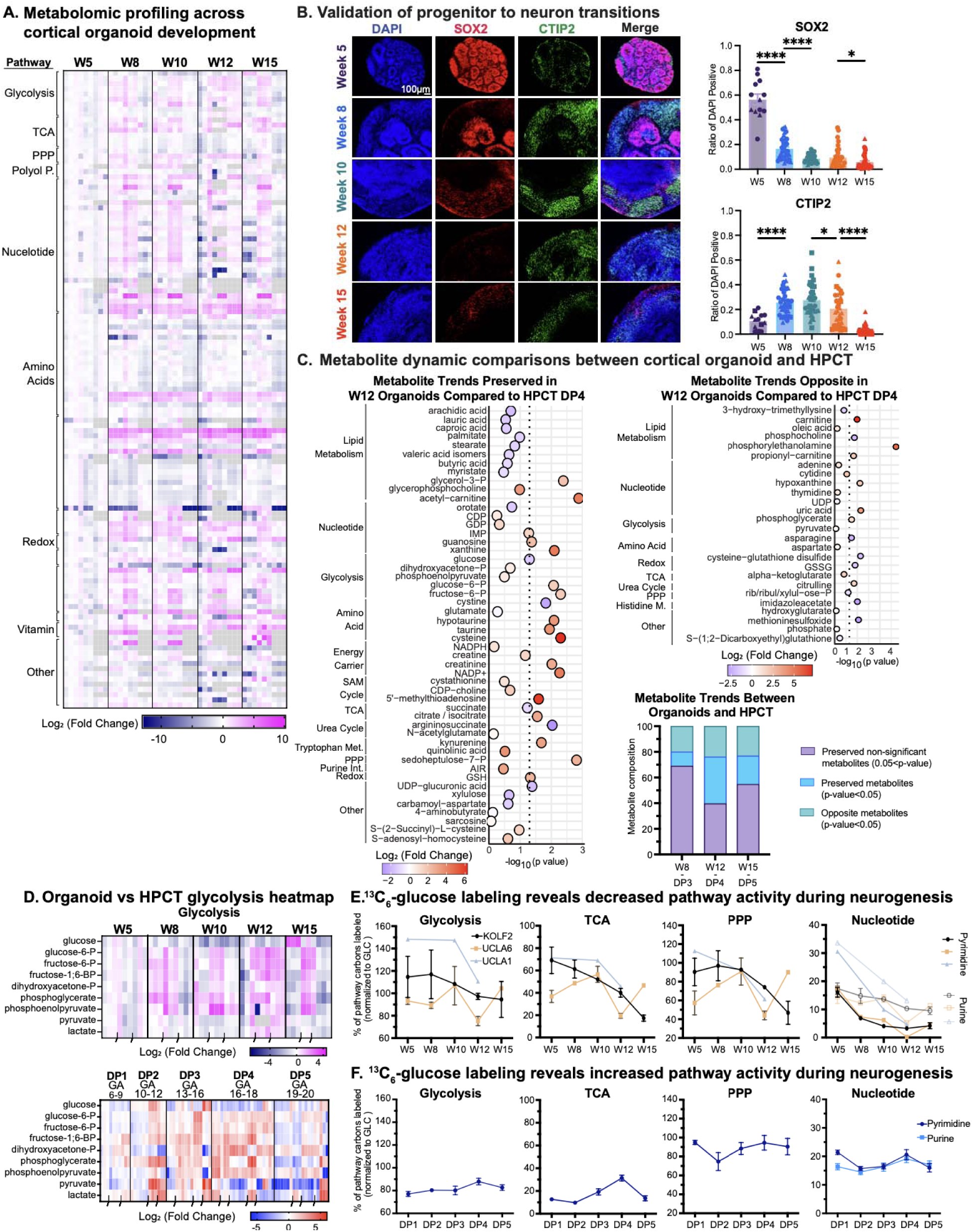
Organoids Recapitulate Key Metabolic Processes of HPCT. A) Cortical organoid metabolic atlas was created using three pluripotent stem cell lines: KOLF2, UCLA6, and UCLA1. Five timepoints of cortical organoid development were measured: week 5 (W5), week 8 (W8), week 10 (W10), week 12 (W12), and week (W15). These five timepoints are known to capture an expansive neurodifferentiation timeline of cortical development while also capturing specific time points of neural differentiation (radial glia emergence, peak neurogenesis, and expected neuronal maturation and gliogenesis). Each time point consisted of three organoid replicates per cell line. Heatmap of 123 targeted metabolite abundances throughout cortical organoid development organized by pathway. Metabolomics was performed on whole organoids using LCMS to measure baseline metabolite levels. Pooled metabolite values were normalized by DNA concentration and TFMS. Each metabolite measurement was scaled to an arbitrary number for data representation. Metabolite levels represent a log_2_ fold change to the youngest sample collected W5. Three biological replicates of each timepoint were analyzed, with three technical replicates for each sample. Blue or pink values indicate a negative or positive log fold change to W5, respectively. Gray values represent an absence of data collected for a metabolite from the metabolomics analysis. Black indents at bottom of heatmap separate biological replicates within each timepoint. B) To validate that cell type composition mirrored expected patterns as in HPCT, we performed IF. A representative image from the KOLF2 line for each of the 5 time points is shown here. Staining for SOX2 (red) was used as a proxy for radial glia progenitor identity, whereas staining for CTIP2 (green) labeled emerging deep layer neuron populations. DAPI, shown in blue, marked nuclei. Scale bar is 100 μM and reflects all images. Quantification of the ratio of DAPI positive cells marked by each protein is presented in bar charts to the right, where 3 distinct biological replicates were analyzed for time point, and 3 technical replicates were used within each biological replicate; shapes on the bar chart correspond to each of the cell lines used (KOLF2 - circles, UCLA6 - square, UCLA1 - triangle). Significance was calculated using the Brown-Forsythe and Welch ANOVA. * indicates p-value < 0.05, ** indicates p-value < 0.01, *** indicates p-value < 0.001. C) Metabolite composition plot showing metabolites trending in the same or opposite direction as in HPCT dataset. Organoid week’s 5, 8, 12, and 15 correspond to HPCT DPs 2, 3, 4, and 5, respectively. Metabolite differential analysis was performed within each system’s dataset (organoid’s metabolite differential analysis compared W8 vs W5, W12 vs W5, and W15 vs W5, while HPCT’s differential analysis compared DP3 vs DP2, DP4 vs DP2, and DP5 vs DP2). We indicate that organoid’s W12 correspond to HPCT’s DP4 and characterize the metabolite trends from each of their differential analyses (through positive or negative log_2_ fold change) as either trending in the same direction, opposite direction, or insignificant metabolite trends in both datasets. Metabolites only needed to be significant in one dataset to be characterized as preserved or opposite metabolites. Metabolites from the organoid’s W12 vs W5 differential analysis that were either preserved or showed opposite trends as the HPCT DP4 vs DP2 are displayed in dot plot. Heatbar displays log_2_ fold change showing trending direction of organoid’s W12 metabolites while x-axis shows whether those metabolites had significance in organoid’s differential analysis. D) Heatmap of glycolytic metabolite levels across cortical organoid (top) and HPCT (bottom) development. Metabolite levels in the heat map represent a log_2_ fold change to each system’s youngest age-HPCT being GA6 and cortical organoid being W5. Three to five biological replicates of each time point were analyzed, with three to six technical replicates for each sample. E) Metabolic pathway tracing graphs of isotopic ^13^C_6_-glucose in glycolysis, TCA, PPP, and nucleotide metabolism across cortical organoid development. Three organoid replicates, per cell line, of each time point were analyzed. Metabolites in glycolysis analysis include G6P, F6P, F16BP, DHAP, 3PG, PEP, and pyruvate. Metabolites in TCA analysis include cis-aconitate, citrate/isocitrate, alpha-ketoglutarate, succinate, fumarate, and malate. Metabolites in PPP include 6PG, RRX5P, and S7P. Metabolites in purine metabolism include hypoxanthine, xanthine, uric acid, AICAR, IMP, inosine, AMP, ADP, ATP, adenine, adenosine, GMP, GDP, GTP, guanine, and guanosine. Metabolites in pyrimidine metabolism include carbamoyl-aspartate, dihydroorotate, orotate, UMP, UDP, UTP, CMP, CDP, CTP, cytidine, and uridine. Y-axis measures the percent of total carbons in pathways that contained ^13^C-label normalized to ^13^C_6_-glucose label; this can result in values over 100% indicating high levels of pathway activity. Pyrimidines and purines in nucleotide metabolism are separated by dark and light colors, respectively (KOLF2 - black circles, UCLA6 - orange squares, UCLA1 - grey triangle). Error bars represent SEM. F) Metabolic pathway tracing graphs of isotopic ^13^C_6_-glucose in glycolysis, TCA, PPP, and nucleotide metabolism across cortical organoid and HCPT development. Three to five biological replicates of each time period were analyzed, with three to six technical replicates for each sample. Metabolites in glycolysis analysis include G6P, F6P, F16BP, DHAP, 3PG, PEP, and pyruvate. Metabolites in TCA analysis include cis-aconitate, citrate/isocitrate, alpha-ketoglutarate, succinate, fumarate, and malate. Metabolites in PPP include 6PG, RRX5P, and S7P. Metabolites in purine metabolism (light blue) include hypoxanthine, xanthine, uric acid, AICAR, IMP, inosine, AMP, ADP, ATP, adenine, adenosine, GMP, GDP, GTP, guanine, and guanosine. Metabolites in pyrimidine metabolism (dark blue) include carbamoyl-aspartate, dihydroorotate, orotate, UMP, UDP, UTP, CMP, CDP, CTP, cytidine, and uridine. Y-axis measures the percent of total carbons in pathways that contained ^13^C-label normalized to ^13^C-glucose label. Error bars represent SEM.

To assess how oxidative stress, a readout of general organoid health, is represented in our samples we analyzed the ratio of glutathione (GSH) over glutathione disulfide (GSSG), which measures the ratio of antioxidants over reactive oxygen species (Zitka et al., 2012). We saw a high ratio in primary samples at DP 2, 3 and 4 with a ratio above 1 at all timepoints, suggesting general cell health. This was matched by very similar dynamics in the organoids (Fig S4B).

### Comparison of Metabolomic Signatures Between Organoids and HPCT Show Preservation of Key Programs

To compare between the datasets with very distinctive provenances, we leveraged a comparison of log_2_fold changes between matching timepoints. Based upon our methylation data, transcriptional analyses (Bhaduri et al., 2020; Camp et al., 2015; Pollen et al., 2019) and biological transitions, the earliest organoid time point (Week 5) matches DP2 in HPCT, so comparisons were performed between organoid weeks 8, 12, and 15 versus week 5 and compared to primary sample ages DP3, DP4, and DP5 versus DP2 (SFig 4C). Pooled analysis showed that ∼80% of metabolites exhibited similar dynamics or were not significantly different between primary tissue and organoids. (Fig 3C). The metabolites trending in the same direction encompassed most pathways represented in the dataset, while a more limited set of pathways had metabolites with opposite trends over developmental time (Fig 3C, SFig 4D & SFig5).

### Differences in Dynamics of Glycolysis Pathway Activity Between Organoids and Primary Samples

Of interest trends in the glycolysis pathway were mostly consistent between datasets, though we noted that almost all of the metabolites in glycolysis intermediates in the pathway before phosphoglycerate follow similar trends (Fig S4E, S5A-C) while the two glycolysis metabolites trending in opposite directions (phosphoglycerate and pyruvate) are found downstream (Fig 3C), suggesting a metabolite specific regulation within this pathway between systems. Consistent with our observations from Seahorse (SFig 3A), zooming into the glycolysis pathway in organoids showed an overall increase in pooled metabolites over developmental time, whereas primary tissue showed a peak at DP4 and diminishing metabolite abundance in DP5 (Fig 3D). These observations are in line with previous reports of increased glycolytic signatures in organoids compared to primary tissue (Bhaduri et al., 2020; He et al., 2024).

In contrast, fractional ^13^C_6_-glucose enrichment analysis showed that the percent of pathway carbons that were labeled in glycolysis were higher in organoids than primary tissue (Fig 3E, SFig 4E), but the pathway activity derived from this tracing analysis showed a decrease over time in organoids only (Fig 3E), while the fractional enrichment analysis matched the pooled tracing in primary tissue with a spike in glycolysis at DP4 (Fig 3E). This suggests that overall, glycolysis is more active in organoids compared to primary tissue, but that movement of carbons through the pathway may be faster at later developmental time points than at earlier time points in cortical organoids (Fig 3E - F). From fractional ^13^C_6_-glucose enrichment analysis, the TCA activity was also generally higher in organoids with a decrease over time, while in HPCT the tracing peaks at DP4. Other pathways adjacent to glycolysis such as the pentose phosphate pathway (PPP) and nucleotide metabolism also had similar trends (Fig 3 E-F, SFig 5A - C). To explore if these pathways were possibly derived from another carbon source, we also included a ^13^C_5_-glutamine label, however we saw that there was minimal labeling (SFig 5D - E) in most pathways.

The metabolomic data showed interesting trends in glycolysis over developmental time, with pieces of the glycolysis pathway being well preserved in organoids. This, paired with ongoing interest in the field about whether glycolysis in organoids impacts cell fate motivated us to directly test if altering glucose in organoid culture could illuminate how glycolysis may influence key cell fate transitions in the context of human cortical development.

### Manipulation of Organoid Media Glucose Changes Cell Types

To determine if glucose manipulations are sufficient to alter glycolytic metabolism in the organoid, and to explore if altered organoid metabolism is sufficient to drive changes in cell fate, we altered organoid media glucose concentrations. Previous studies have shown that neural stem cells (NSCs) typically have higher glycolysis, with shifts towards OXPHOS with differentiation (Traxler et al., 2021; Zheng et al., 2016). Moreover, lowered environmental glucose in stem cell media conditions has been shown to drive maturation in other tissues including cardiac muscle (Nakano et al., 2017). However, in the context of human cortical development, the links between glucose, differentiation and cell type identity, and a connection to metabolite composition have not been thoroughly explored.

Thus, we chronically exposed Week 5 (W5) organoids, which are primarily radial glia enriched, to three glucose concentrations in the media until week 12 (W12) of organoid development. The three concentrations were 17.5 mM (standard media glucose), 8.75 mM (reduced glucose), and 2.5 mM glucose, which has been reported to be the physiological level in the brain (Kleman et al., 2008). To analyze the experiment, we performed the same suite of analyses as done previously, this time also including single-nucleus RNA-sequencing (snRNA-seq) as an additional method to characterize broad changes in cell types and to identify impacted gene programs (Fig 4A). Immunofluorescence analysis of lowered media glucose conditions showed a significant decrease in deep layer neurons, while astrocytes (marked by GFAP) and oRG populations (marked by HOPX) increased significantly across three organoid lines (Fig 4B). Increased expression of cleaved-caspase 3 (CC3) was observed in the center of the organoid across all our conditions as they increase in age and size, which is expected for a W12 cortical organoid. However, we also see evidence of a greater necrotic core in organoids cultured in lower glucose levels (2.5mM) (SFig 6A).

**Figure 4:**
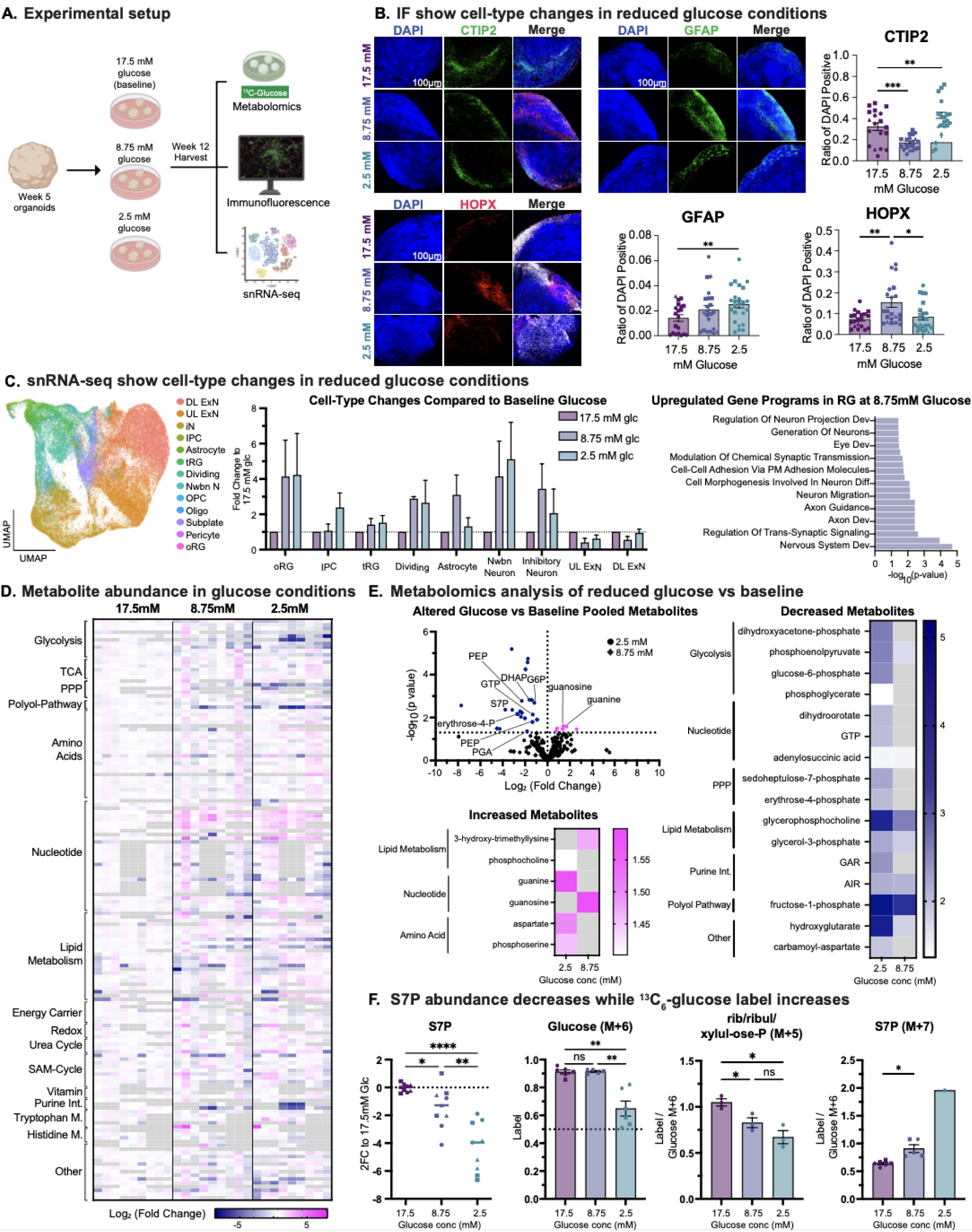
Glucose Manipulation Alters Organoid Cell Types and Pentose Phosphate Pathway. A) Schematic of cortical organoids undergoing glucose manipulation processing pipeline. Cortical organoids were cultured for 5 weeks and then divided into three groups: cultured in 17.5mM glucose levels (control group), 8.75mM glucose, and 2.5mM glucose. Cortical organoids were cultured in their respective glucose level for 7 weeks and harvested at their week 12 time point for metabolomics and isotopic ^13^C_6_-glucose tracing, immunofluorescence to evaluate cell type composition, and single-nuclei RNA sequencing (snRNA-seq) to further characterize cell types and genetic programs that were changed in altered glucose conditions. B) To validate that cell types, change in altered glucose conditions, we performed immunofluorescence. A representative image from the KOLF2 line for each of the 3 condition groups is shown here. Staining for CTIP2 (green) labeled emerging deep layer neuron populations, staining for GFAP (green) labeled astrocyte populations, and staining for HOPX (red) was used as a proxy for outer-radial glia (oRG) progenitor identity. DAPI, shown in blue, marked nuclei. Scale bar is 100 μM and reflects all images. Quantification of the percent of DAPI positive cells marked by each protein is presented in bar charts to the right, where three lines of organoids were analyzed for each condition, and three organoid replicates per cell line were used; shapes on the bar chart correspond to each of the cell lines used (KOLF2 - circles, UCLA6 - square, UCLA1 - triangle). Significance was calculated using the Brown-Forsythe and Welch ANOVA. * indicates p-value < 0.05, ** indicates p-value < 0.01, *** indicates p-value < 0.001. C) snRNA-seq analysis was performed at week 12 across three organoid lines and all three glucose concentration conditions. Left: UMAP shown here represents all 81,593 cells passing quality control across lines and conditions. Cell types were annotated by mapping to an atlas of human cortical development (Nano et al., 2023) and UMAP is colored by these annotations encompassing expected cell populations. Middle: Cell type fold changes were calculated by taking the fraction of cells in each glucose concentration and comparing it to the paired fraction in the baseline (17.5 mM) condition. These data highlight an increase in progenitor and neuronal populations. Right: Gene Ontology Biological Process analysis of the genes upregulated in radial glia (RG) at the 8.75 mM condition highlight gene programs related to neuronal development. D) Heatmap of 155 targeted metabolite abundances across three glucose level conditions-17.5mM (control level), 8.75mM, and 2.5mM. LCMS-based metabolomics was performed on whole organoids to measure baseline metabolite levels. Pooled metabolite values were normalized by DNA concentration and trifluoromethanesulfonate (TFMS). Each metabolite measurement was scaled to an arbitrary number for data representation. Metabolite levels represent a log_2_ fold change to the baseline/control glucose level (17.5mM). Each condition group consisted of three cell lines with three organoid replicates per cell line. Dark blue or pink values indicate a negative or positive log_2_ fold change to 17.5mM glucose, respectively. Gray values represent an absence of data collected for a metabolite from the metabolomics analysis. Black indents at bottom of heatmap separate biological replicates within each condition group. E) Volcano plot highlighting differential metabolite analysis between altered glucose conditions (8.75mM-diamond and 2.5mM-circle) compared to control/baseline glucose (17.5mM). The X-axis shows log_2_(fold change) of altered glucose metabolites compared to the baseline glucose group. The Y-axis shows -log_10_(P-Value) of each metabolite between condition groups with a dashed line at the 1.3 mark to highlight metabolites that were significant. Breakdown of significantly increased metabolites with their corresponding pathways are shown in the heatbar with pink heatbar legend displaying its respective -log_10_(P-Value). Breakdown of significantly decreased metabolites with their corresponding pathways are shown in the heatbar with blue heatbar legend displaying its respective -log_10_(P-Value). F) Sedoheptulose-7-Phosphate (S7P) was a particular metabolite of interest. Dot plot of S7P abundance log_2_ fold change in altered glucose conditions compared to control/baseline glucose (17.5mM) is shown. Additionally, isotopic ^13^C-label derived from ^13^C_6_-glucose in Glucose M+6, RRXP M+5, and S7P M+7 are plotted by condition groups. Each condition group consisted of three cell lines with three organoid replicates per cell line; shapes on the bar chart correspond to each of the cell lines used (KOLF2 - circles, UCLA6 - square, UCLA1 - triangle). Significance was calculated using the Brown-Forsythe and Welch ANOVA. * indicates p-value < 0.05, ** indicates p-value < 0.01, *** indicates p-value < 0.001.

To link these cell type changes to transcriptional identity, we profiled all three lines across the three media conditions using snRNA-seq, resulting in data from 81,593 high quality cells (STable 7). We performed standard quality control analysis and mapped cell types in our dataset using a previously annotated meta-atlas (Nano et al., 2023) of human cortical development (Fig 4C). We observed how our data were distributed across lines, treatments, and verified by marker gene expression (SFig 6B - D). When we explored composition across each treatment in each of the three lines, we found similar changes in cell type to what was observed in the IF analysis. Namely, astrocytes and oRGs increased at least one of the lowered glucose concentrations; additionally, we saw an increase in intermediate progenitor cells (IPCs), dividing cells, newborn neurons and inhibitory neurons (Fig 4C). Interestingly, these changes in cell type were specific to individual cell types and lineages, as opposed to a uniform change across the whole dataset. To interrogate whether these changes were due to changes in maturation or due to underlying differences amongst the neural progenitors, we performed differential gene expression across conditions. Gene ontology (GO) terms show significant changes in RG with increased gene programs related to neuronal development and synaptic biology at the 8.75 mM condition, indicating this could be a cell fate switch in RG. We also observed that gene modules related to glycolysis and PPP were significantly downregulated with reduced glucose, while the TCA cycle genes were significantly increased (SFig 6E).

To find the metabolic mechanisms underlying this change in cell types, we use LCMS-based metabolomics and ^13^C_6_-glucose tracing (STable 8 - 10). Across 155 metabolites we saw broad shifts in metabolic programs across the three lines, with changes in pathways linked to glycolytic metabolism (Fig 4D). This indicates that although this was manipulation of a key nutrient metabolite, the metabolism effects were largely localized to expected pathways (Fig 4E). Differential metabolite analysis of both reduced glucose conditions compared to 17.5 mM baseline glucose showed numerous significantly changing metabolites, with most being downregulated (Fig 4E, SFig 6F-H). Overall, we observed greater magnitude decreases in metabolite levels in lowered glucose conditions, and often in metabolites from branch pathways off of glycolysis, including the PPP, polyol pathway, and nucleotide intermediates (Fig 4E, SFig 6G). We observed notable increases in guanine, guanosine, aspartate, phosphoserine, 3-hydroxy-trimethyllysine, and phosphocholine (Fig 4E, SFig 6F). The redox cycle branching from the PPP was largely unaffected by decreases in glucose concentrations indicating similar levels of cell health (SFig 6H).

We further explored the PPP and observed significantly decreased levels of sedoheptulose-7-phosphate (S7P), a central metabolite in this pathway, in the lowered glucose concentrations (Fig 4F). This was consistent with our observations of decreased ^13^C_6_-glucose labeling of other relevant PPP metabolites including rib/ribul/xylul-ose-P (RRXP) in lowered glucose concentrations. However, we observed increased ^13^C_6_-glucose-derived labelled S7P in lowered glucose concentrations, suggesting a potentially important role for S7P and the PPP in the cell-type transitions observed with decreased glucose.

### PPP Pathway Exhibits Temporally Specific Dynamics Across Systems of Human Cortical Development

Our observations from the glucose manipulation experiment drew our attention to the PPP. The PPP is comprised of two major branches (Stincone et al., 2015): the oxidative branch generates NADPH which is a critical reducing agent for many biosynthetic reactions as well as the GSH/GSSG redox cycle (Ge et al., 2020), while the non-oxidative branch feeds into nucleotide synthesis, often driving the requirement for nucleotides in highly proliferative cells (Fig 5A). In cardiac muscle, high glucose has been shown to impair maturation through excessive promotion of nucleotide biosynthesis mediated by the PPP (Nakano et al., 2017). Our data raised the possibility that the PPP may be also related to cell fate specification decisions made by RG through human cortical development. Thus, we explored the dynamics of PPP metabolites in our atlas data.

**Figure 5:**
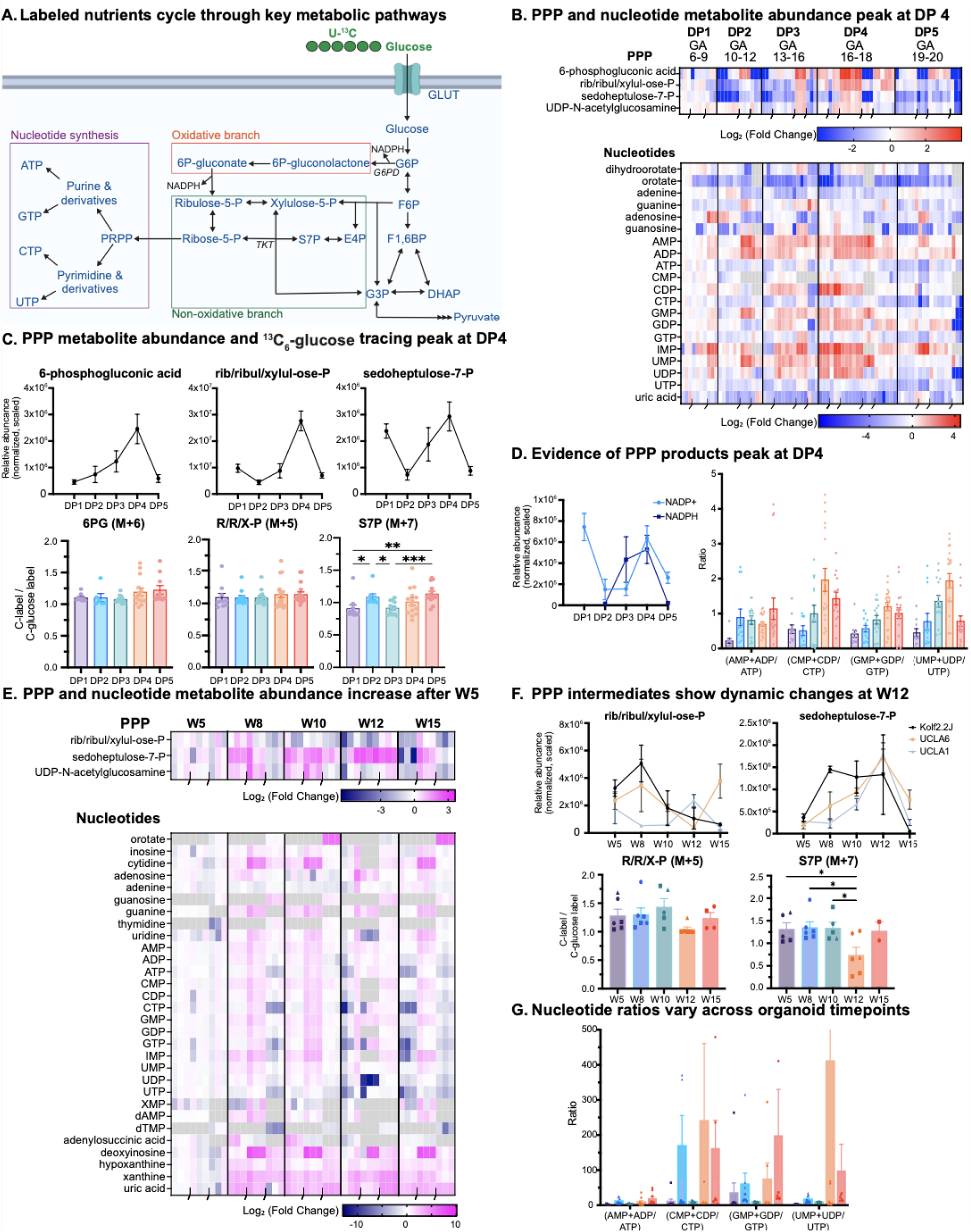
PPP Dynamics are Preserved Between HPCT and Organoid Mimicking Glycolysis. A) Schematic of glycolysis and pentose phosphate pathway (PPP). Glycolysis begins with glucose entering the cell from the cytoplasm. The PPP pathway consists of two arms: oxidative branch (boxed in green) and the non-oxidative branch (boxed in orange). The oxidative branch is known for its production of NADPH, while the non-oxidative branch contributes to nucleotide synthesis (boxed in purple). B) HPCT heatmaps of PPP and nucleotide related metabolite levels peak at DP4. Metabolite levels in the heatmap represent a log_2_ fold change to GA6. Three to five biological replicates of each DP were analyzed, with three to six technical replicates for each sample. C) Metabolite abundance graphs and ^13^C_6_-glucose label of 6-phosphogluconic acid (6PG), rib/ribul/xylul-ose-phosphate (RRXP), and S7P validates pooled metabolomics showing a peak at DP4 for PPP metabolites. Three to five biological replicates of each DP were analyzed, with three to six technical replicates for each sample. Combined metabolite abundance values were calculated by averaging all technical replicates across biological samples within each DP. Glucose tracing graphs display fully labeled isotopologues indicated by M+5/6/7. The y-axis measures abundance (measured and scaled) across each DP or ratio of ^13^C-label normalized to ^13^C_6_-glucose label (M+6). Dots show technical replicates from each biological sample within DPs. Error bars represent SEM. D) To further analyze PPP activity, we looked at products of PPP. Left graph is the metabolite abundance of NADP+, an essential precursor metabolite, and NADPH, a key oxidative branch product. Right graph shows the ratio of mono- and di-phosphate over tri-phosphate metabolite abundance of adenosine, cytidine, guanosine, and uridine nucleotides to evaluate nucleotide production as an indicator of the non-oxidative branch activity. Both PPP products show peaks at DP4 further validating pooled metabolomics showing a peak of PPP pathway at DP4. Three to five biological replicates of each DP were analyzed, with three to six technical replicates for each sample. Combined metabolite abundance values were calculated by averaging all technical replicates across biological samples within each DP. Dots show technical replicates from each biological sample within DPs. Error bars represent SEM. E) Cortical organoid heatmaps of PPP and nucleotide related metabolites across five time points. Metabolite levels in the heatmap represent a log_2_ fold change to W5. Each time point consisted of three cell lines with three organoid replicates per cell line. F) Metabolite abundance graphs and ^13^C_6_-glucose label of RRXP and S7P in cortical organoid development across five time points. Each time point consisted of three cell lines with three organoid replicates per cell line. Combined metabolite abundance values were calculated by averaging all technical replicates across organoid samples within each time point. Glucose tracing graphs display fully labeled isotopologues indicated by M+5/6/7. The y-axis measures abundance (measured and scaled) across each DP or ratio of ^13^C-label normalized to ^13^C_6_-glucose label (M+6) labeling. Dots show technical replicates from each biological sample within DPs. Error bars represent SEM. G) Mono- and di-phosphate over tri-phosphate nucleotide ratios metabolite abundance of adenosine, cytidine, guanosine, and uridine nucleotides to evaluate nucleotide production as an indicator of the non-oxidative branch activity. Each time point consisted of three cell lines with three organoid replicates per cell line. Dots show technical replicates from each biological sample within DPs. Error bars represent SEM.

When looking at our HPCT metabolic atlas, the PPP, like glycolysis, peaks at DP4 (Fig 5B). This is seen across individual metabolites when analyzing the changes in their abundance over time which each also peak at DP4. The ^13^C-label fully labeled isotopologues of 6-phoshogluconic acid (GP6), RRXP, and S7P remain stable and highly labeled over time (Fig 5C). We additionally observed that products of the PPP pathway, including NADPH peaks at DP4 as does the NADPH/NADP+ ratio which is indicative of activity in the oxidative branch of the PPP (SFig 7A). We also looked into downstream activity of PPP by quantifying the abundance of mono- and di-phosphate nucleotides over the tri-phosphate between purines and pyrimidines. We see an overall increase in mono- and di-phosphate over tri-phosphate adenosine nucleotides, but minimal changes in guanosine ratio in HPCT (SFig 7B). When looking at ^13^C_6_-glucose tracing, we saw a general increase of ^13^C-label in adenosine over time, but a subtle decrease in guanosine (SFig 7C). We noted an increase in mono- and di-phosphate over triphosphate ratios for pyrimidines (SFig 7D). The ^13^C-label showed dynamic changes over time in pyrimidines (SFig 7E). We applied the same analysis to the organoid dataset and saw an increase in mono- and di-phosphate over tri-phosphate purine ratios, but a general decrease in ^13^C-label in these nucleotides (SFig 7F, 7G). We see a similar trend in mono- and di-phosphate over tri-phosphate ratios and ^13^C-label in pyrimidines (SFig 7H, 7I). Together these data, in line with the increase in glycolysis and the PPP in DP4, suggest that there is a high prioritization of carbons from glucose being resourced towards nucleotide synthesis at this specific developmental time period, with the PPP as a crucial intermediate pathway.

We explored our organoid metabolic atlas and similarly saw an increase in PPP intermediates and nucleotides at matching timepoints based upon metabolomics (Fig 5E). Additionally, S7P abundance spiked at week 12, though its ^13^C-label decreased at this timepoint, opposite of what we see in our glucose depletion experiment (Fig 5F). Parallel dynamics of nucleotide ratios in the organoid compared to primary were observed (Fig 5G, SFig 7C or D nucleotide dynamics). Together, these data showed that general PPP trends were preserved between the primary and organoid. Given our previous glucose experiment showing interesting S7P dynamics, the matched timing of the PPP and glycolysis peaking in DP4 in primary samples, and the preservation of upstream glycolysis metabolites in organoids and primary tissue, we sought to experimentally modulate PPP and test the hypothesis that it is a regulator of cell fate specification in human cortical development.

### Pharmacologic Inhibition of PPP Phenocopies Key Features of Glucose Manipulations

To functionally test the impact of PPP on cell fate specification and metabolism, we pharmacologically inhibited glucose-6-phosphate dehydrogenase (G6PD) and transketolase (TKT). We administered the inhibitors at week 8 (W8) to mimic the stages before the PPP peaks in the organoid and matched timepoints in primary DP4. The two inhibitors, 6-Aminonicotinamide (6AN) targeting G6PD in the oxidative branch, and oroxylin A (ORX) targeting TKT’s bidirectional activity in the non-oxidative branch, were used at concentrations from published studies (Jia et al., 2022; Nakano et al., 2017). The inhibitors were delivered to three cell lines and maintained in the media for 4 weeks, with cells harvested at week 12 (W12) for metabolomics, IF, and snRNA-seq. Analysis of the cell types with IF showed a significant increase in mature neurons, astrocytes, and outer radial glia cells in drug versus DMSO control conditions (Fig 6B, SFig 8A).

**Figure 6:**
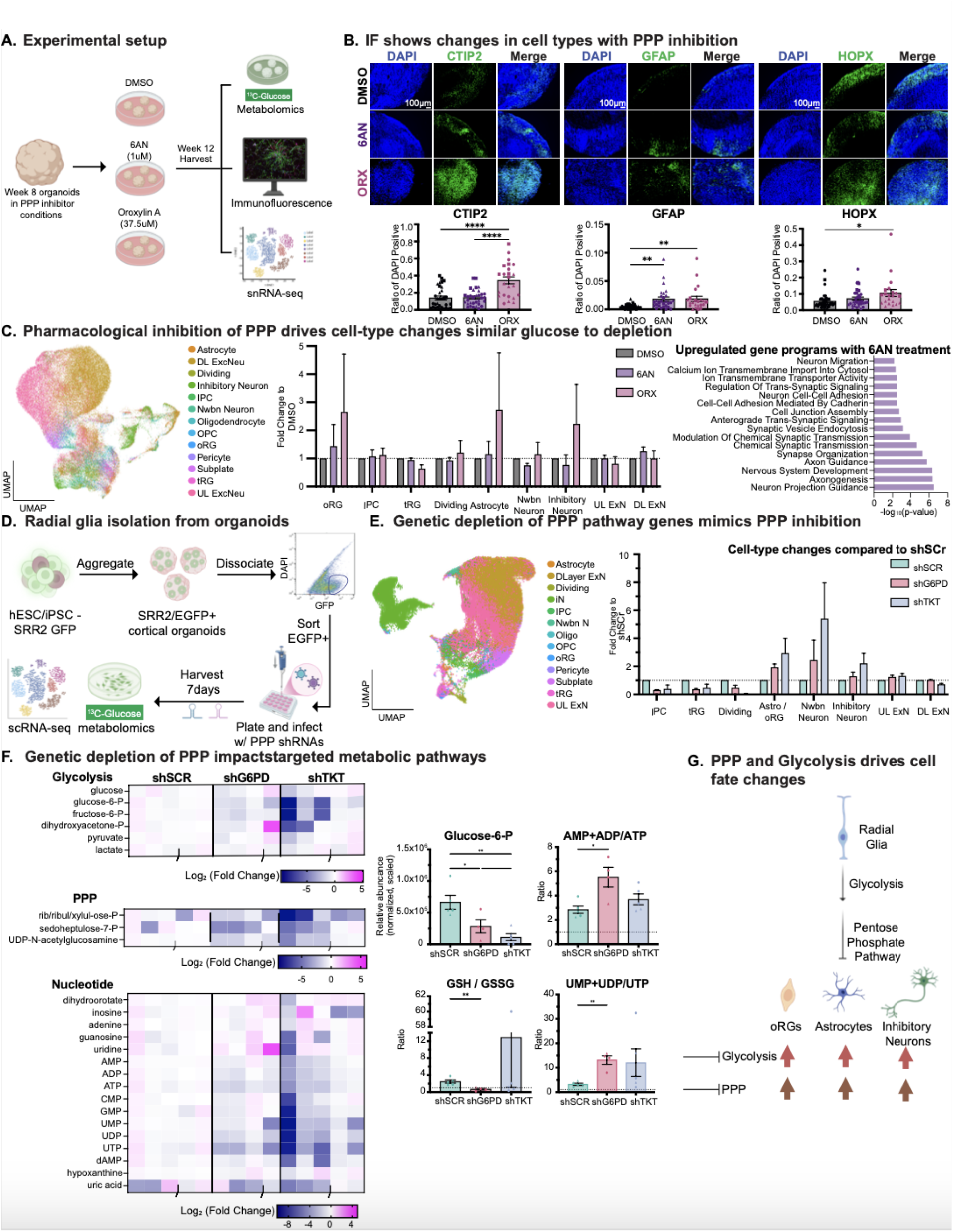
PPP Pathway Modulates Radial Glia Cell Fate Specification. A) Schematic of cortical organoids undergoing pharmacological inhibition of PPP (PPPi) enzymes processing pipeline. Cortical organoids were cultured for 8 weeks and then divided into 3 groups: cultured with DMSO (control group), 6-Aminonicotinamide (6AN) at 1uM, and Oroxylin A (ORX) at 37.5uM. Cortical organoids were cultured in their respective conditions for 4 weeks and harvested at their W12 time point for metabolomics and ^13^C_6_-glucose tracing, immunofluorescence to evaluate cell type composition, and snRNA-seq to further characterize cell types and genetic programs that were changed in PPPi manipulations. B) To validate that cell type changes with pharmacological inhibition of PPP, we performed immunofluorescence. A representative image from the KOLF2 line for each of the three condition groups is shown here. Staining for CTIP2 (green) labeled emerging deep layer neuron populations, staining for GFAP (green) labeled astrocyte populations, and staining for HOPX (green) was used as a proxy for oRG progenitor identity. DAPI, shown in blue, marked nuclei. Scale bar is 100 μM and reflects all images. Quantification of the percent of DAPI positive cells marked by each protein is presented in bar charts to the right, where three cell lines and three organoid replicates per cell line were analyzed for each condition; shapes on the bar chart correspond to each of the cell lines used (KOLF2 - circles, UCLA6 - square, UCLA1 - triangle). Significance was calculated using the Brown-Forsythe and Welch ANOVA. * indicates p-value < 0.05, ** indicates p-value < 0.01, *** indicates p-value < 0.001. C) snRNA-seq analysis was performed at Week 12 across three organoid lines and all three PPP pharmacologic inhibitor conditions. Left: UMAP shown here represents all 106,047 cells passing quality control across lines and conditions. Cell types were annotated by mapping to an atlas of human cortical development (Nano et al., 2023) and UMAP is colored by these annotations encompassing expected cell populations. Middle: Cell type fold changes were calculated by taking the fraction of cells in each treatment condition and comparing to the paired fraction in the vehicle only (DMSO) condition. These data highlight a phenocopy of the glucose depletion experiment with increased oRG, astrocyte, and inhibitory neuron populations. Right: Gene Ontology Biological Process analysis of the genes upregulated in the 6AN treated cells inhibiting G6PD highlight gene programs related to neuronal development. D) Schematic of workflow to isolate radial glia from organoids and knockdown PPP enzymes using shRNA. hPSC lines were infected with lentivirus with SRR2, an enhancer for SOX2 which is highly expressed in radial glia, regulating expression of EGFP as a fluorescent reporter. hPSC SRR2/EGFP stem cells were used to generate cortical organoids. Beginning at week 8, organoids were dissociated, and radial glia were then isolated through FACS of GFP+ cells. Cells were immediately plated and infected with shRNA constructs targeting G6PD, an enzyme in the oxidative branch, and TKT, an enzyme in the non-oxidative branch. Cells were cultured for 7 days and then harvested for metabolomics and scRNA-seq analysis. E) scRNA-seq analysis was performed 1 week after radial glia enrichment from W8 organoids and treatment with knockdown hairpins. Left: UMAP shown here represents all 51,105 cells passing quality control across lines and conditions. Cell types were annotated by mapping to an atlas of human cortical development (Nano et al., 2023) and UMAP is colored by these annotations encompassing expected cell populations. Right: Cell type fold changes were calculated by taking the fraction of cells in each knockdown and comparing to the paired fraction in the scrambled control. These data highlight a phenocopy of the glucose depletion and pharmacological PPP inhibition experiment with increased astrocytes/outer radial glia and inhibitory neuron populations. F) Cortical organoid heatmaps of glycolysis, PPP, and nucleotide related metabolites across treatment with knockdown hairpins. Metabolite levels in the heatmap represent a log_2_ fold change to short-hairpin RNA scramble (shSCR) control. Additionally, we measure metabolite abundance of glucose-6-phosphate (G6P) across treatment with knockdown hairpins. Oxidative stress levels are shown in the GSH/GSSG ratio plot. We also measured mono- and di-phosphate over tri-phosphate nucleotide ratios metabolite abundance of adenosine and uridine nucleotides to evaluate nucleotide production as an indicator of the non-oxidative branch activity. Each time point consisted of three cell lines with three organoid replicates. Combined metabolite abundance values were calculated by averaging all technical replicates across organoid samples within each time point. Dots show technical replicates from each biological sample within DPs. Error bars represent SEM. Significance was calculated using the Brown-Forsythe and Welch ANOVA. * indicates p-value < 0.05, ** indicates p-value < 0.01, *** indicates p-value < 0.001. G) Schematic shows glycolysis feeding into PPP and regulating cell fate decisions by restricting radial glia from becoming oRGs, astrocytes, and inhibitory neurons prematurely in development. Using experimental manipulations, we observed that inhibiting glycolysis and PPP resulted in increased fractions of these 3 cell types.

snRNA-seq of all three lines and three treatment conditions yielded 106,047 high quality cells and were again analyzed for distributions of lines, treatment conditions, and validated for expected marker gene composition (SFig 8B, STable 11). The cell types were annotated as previously, and showed dynamics that paralleled cell type changes in depleted glucose conditions (Fig 4B); notably, this was stronger with the ORX that targets TKT compared to the 6AN targeting G6PD (Fig 6C, SFig 8B-D). However, we did observe that 6AN treatment significantly decreased the expression of PPP genes (SFig 8E) and drove changes in genes related to neuronal differentiation, similar to our glucose depletion conditions (Fig 4B).

Metabolomic analysis of the organoids at W12 showed limited changes across conditions in most pathways, including the PPP (SFig 8F, STables 12 - 14). We hypothesized that this may be indicative of the organoid cells adjusting to inhibitor exposure and rewiring their cellular metabolism which is a well-described phenomenon when drugs inhibiting metabolic enzymes are used (Zaal and Berkers, 2018), thus mitigating more obvious impacts upon the PPP pathway. However, because the cell type changes closely phenocopied what we observed in the glucose depletion experiment, this suggested that inhibition of the PPP either drives a change in fitness or directly impacts the cell fate choices being made by cortical RG.

### Genetic Inhibition of PPP Links Metabolism to Radial Glia Cell Fate

To explore the exciting possibility that the PPP impacts RG cell fate specification, we performed an experiment where we used a reporter for SOX2 activity driven by an enhancer known as SRR2 (SOX2 regulatory region 2) that has been described to specifically drive SOX2 expression (Sikorska et al., 2008). This enhancer has also been shown to be selectively active in radial glia in two independent single-cell/nuclei epigenomics datasets of the developing human brain (Wang et al., 2025; Ziffra et al., 2021), suggesting that we could specifically isolate radial glia by using an SRR2 reporter. We therefore grew organoids with the SRR2/EGFP and validated that the SRR2/EGFP+ positive cells were SOX2 positive RG (SFig 9A). We then isolated these cells from the organoid using fluorescent activated cell sorting (FACS) and plated them in the presence of a lentivirus delivering a short-hairpin RNA (shRNA) targeting *G6PD* or *TKT,* which we observed robust knockdown via real-time PCR (Fig 6D, SFig 9B). By validating that the cells that were purified were indeed SOX2 positive (SFig 9C (plots), SFig 9D (intracellular FACS) and largely depleted of CTIP2, a neuronal marker (SFig 9D (intracellular FACS)), we tested how knockdown of these PPP genes might impact RG cell fate and metabolism.

A previous study that characterized a related TKT gene, hTKTL1 showed that PPP activity is important in driving oRG expansion in the human cortex, which previews an important role for this pathway in modulating oRG. However, our knockdown targets a distinct version of this gene (TKT) alongside G6PD at a specific time point during which PPP was highly active from our metabolic atlas, which differs from the experimental design of their study (Pinson et al., 2022).

After 1 week in culture, we performed single-cell RNA-sequencing (scRNA-seq) to characterize the cell types across the three stem cell lines that we analyzed (Fig 6E, SFig 10A-C). This resulted in high quality data from 51,105 cells that were mapped to our meta-atlas; we also explored how these cell types were distributed across lines and knockdown conditions (STable 15, SFig 10A-C). Intriguingly, we observed some of the same cell type changes, specifically in our oRG/astrocyte cluster, newborn neurons, and inhibitory populations (Fig 6E). PPP inhibition affected a subset of cells identified in the glucose depletion experiment, specifically shifting radial glia toward a more mature population that generates neurons, astrocytes, and a recently described dorsal-derived inhibitory neuron lineage in HPCT and organoids (Andrews et al., 2023; Delgado et al., 2022). These shifts also mimic the predicted daughter cells of a recently described Tri-IPC progenitor in human development (Wang et al., 2025). The PPP module genes decreased significantly with TKT knockdown but corresponded to an increase in the nucleotide gene program analyzed (SFig 10D).

Our metabolomics analysis showed strong and sustained changes in PPP and nucleotide metabolite abundance, with an unexpected feedback loop that drove a decrease in glycolysis metabolite abundance as well (Fig 6F). We see evidence of decreased glucose-6-phosphate (G6P) pool abundance in both shG6PD and shTKT, giving additional evidence of a feedback loop from the knockdown of these PPP enzymes (Fig 6F-G). However, we also see increases in certain PPP indirect products, such as GSH in which we see an increase in GSH / GSSG ratio in the shTKT condition. In addition, we also noted increases in mono- and di-phosphate nucleotides, specifically in adenosine and uridine nucleotides, in the shG6PD condition (Fig 6G). This was also true when analyzing the ^13^C_6_-glucose labelled metabolites where the shTKT had dramatic impacts on nucleotide tracing, with a decrease of these same labelled fractions in the cells with shG6PD as well (SFig 9E).

Together, these results propose a model where the PPP, fed by glycolysis, is essential for the suppression of radial glia maturation and generation of some, but not all, daughter cell types. We specifically observe that glycolysis and PPP peak in activity at DP4, and that this peak is matched for PPP in cortical organoid systems. When PPP is inhibited indirectly through reduced glucose, or directly through pharmacologic inhibition or genetic knockdown, we observe an increase in oRGs, astrocytes, newborn neurons, and inhibitory neurons (Fig 6H). This suggests a key role for the PPP in human cortical cell fate specification and provides a dataset from which additional pathways can be experimentally interrogated in this context.

## Discussion

### Cell Fate Changes Mediated by Glycolysis Through PPP

Here, we leverage our atlas of metabolism in the developing human cortex using both HPCT and organoid samples to discover a temporally specific role for glycolysis and the PPP to regulate cell fate specification. Through extracellular depletion of glucose, we observed cell type compositional changes in the organoid that were recapitulated when we inhibited PPP activity through both pharmacological and genetic inhibition, highlighting that intracellular manipulation of PPP also impacted cell fate decisions during cortical development. Across experiments, we observe a common shift in increasing outer radial glia cells, inhibitory neurons, and astrocytes. This suggests that PPP activity may serve as an inhibitor of radial glia maturation where it prevents the premature transition to outer radial glia that then go on to become gliogenic. Consistent with this shift in maturation state, the changes in cell fate specification such as the generation of inhibitory neurons is consistent with a recently described late stage phenomena of radial glia dorsally producing inhibitory neurons (Andrews et al., 2023; Delgado et al., 2022).

While astrocytes are known to rely on glycolytic activity as their main energy source (Dienel and Hertz, 2001), we see that depleting glucose and downregulating PPP shows an increase in astrocyte populations. Our study shows counterintuitive results to the established metabolic profile of astrocytes and therefore may further support the scenario of cell fate changes occurring rather than a metabolic remodeling for survival. In the context of these cell fate changes, we were intrigued that both the cell type shifts and the transcriptional programs impacted by altered external glucose levels or internal PPP modulation occurred in a highly cell type-specific manner, with individual cell types such as radial glia experiencing substantial changes in transcriptional programs while other cell types, including many upper and deep layer neurons, remained largely unaffected.

The changes in cell type with the modulations we performed in this study, such as the increase in oRG cells with lowered glucose concentrations, have long been speculated to be necessary changes to improve the fidelity of organoids to primary tissue at comparable stages. However, our studies here suggest that there is not a fully linear relationship between lowered glucose and improved fidelity to cell types in human cortical development. For example, some features such as the increase in astrocyte populations earlier, may be helpful in modeling their growth, but we also observed additional cell death in the necrotic core in decreased glucose conditions, suggesting that some of the changes in cell type were mitigated by poorer organoid health. Additional investigation of how metabolism and culture conditions drive the underlying biology of cell fate specification, maintenance, and maturation will enable more nuanced improvements to organoid culture conditions in the future.

### Links Between Metabolism and Cell Fate

Based upon these observations and cell type-specific changes, we identify a role for the PPP in modulating cell fate specification in the developing human cortex. In previous studies, metabolites have been shown to play a role in driving shifts in differentiation and maturation, though independently many of these studies have converged on lactate and its related enzymes as modulators of differentiation and maturation. The PPP has been associated with inhibiting maturation in cardiomyocytes based upon promotion of nucleotide biosynthesis (Nakano et al., 2017). While our data also shows changes in nucleotide levels and ratios, we additionally observe a potentially central role of S7P as a molecule whose levels appear to be carefully constrained by the cell when subjected to altered glucose levels or PPP activity. These data suggest that PPP may play a dynamic role in regulating neural stem cell maturation and its downstream choice of cell fate.

The links between PPP and cell fate evoke questions as to whether alterations of this pathway could be implicated in neurodevelopmental disorders. Indeed, the PPP is known to regulate neuroinflammation (Tu et al., 2019), a key risk factor for autism spectrum disorder (Matta et al., 2019). Additionally, G6PD deficiencies have been linked to individual patients with epilepsy (Liguori et al., 2013; Merdin et al., 2014) and ribose-5-isomerase deficiency syndrome is an inborn error of metabolism with developmental delay as a symptom (Huck et al., 2004). These examples, paired with our observations, suggest that altered PPP intermediates could be a mechanism underlying neurodevelopmental disorders more broadly.

### Resource for the Field

Our ability to leverage this atlas to discover and investigate the temporal importance of PPP metabolism at DP4 shows that this data set could yield novel inquiries into other metabolic pathways and their role in cortical development. To this end, we have provided all raw and processed tables for others to further explore our metabolomics atlas and to derive additional hypotheses. Moreover, data from both HPCT and cortical organoids informed us of preserved metabolic pathways between systems, and our manipulation of glycolysis and PPP paired with transcriptional characterization allowed us to begin linking changes in metabolic properties to RNA-based cell type changes, providing a unique multiomic perspective to our data. Future efforts will expand on this resource and continue to more completely dissect the ways in which metabolism drives individual cell type processes during development.

## Acknowledgments

We would like to thank the members of the Bhaduri and the Christofk Labs (especially E. Fazzari, S. Campbell and B. Wilde) for their insightful advice and comments on the study. We would like to thank the Broad Stem Cell Research Center Imaging Core, Suhua Feng (UCLA Broad Stem Cell Research Center Sequencing Core) for help with running sequencing. Thank you to Roxane Verdikt and Patrick Allard for media resources, and to Ko Kiehle for tissue procurement. The work performed in the manuscript was generously funded by R00NS111731 from the NIH (NINDS), R01MH132689 from the NIH (NIMH), the Young Investigator Award from the Brain & Behavior Research Foundation, the Alfred P. Sloan Foundation, the Rose Hills Foundation, the Klingenstein-Simons Fellowship from the Esther A. & Joseph Klingenstein Fund and the Simons Foundation, the NIH BRAIN Initiative Cell Atlas Network (UM1MH130991), the Chan Zuckerberg Initiative (to A.B. and H.R.C), and Ablon Scholar Award (to A.B.). Additional funding was provided to P.R.N. and J.M. (UCLA Eli and Edythe Broad Center of Regenerative Medicine and Stem Cell Research Training Program), C.V.N. (T32 NS048004, Predoctoral Fellowship in association with the Training Grant in Neurobehavioral Genetics), and R.L.K. (T32 GM145388, Cell and Molecular Biology Training Program). This material is based upon work supported by the National Science Foundation Graduate Research Fellowship Program under Grant No. (DGE-2034835) to J.S. Any opinions, findings, and conclusions or recommendations expressed in this material are those of the author(s) and do not necessarily reflect the views of the National Science Foundation. Funding was also received from the Eugene V. Cota-Robles Award to J.S.

## Author Contributions

Conceptualization - AB ; Methodology - JM and JS; Validation - JS and JM; Formal Analysis - JM, JS, AB, NM, AK, FD, LS, KPM, PRN, AAM; Investigation - JM, JS, NM, AK, LS, KPM, DJA, CEG, PRN, AAM, CAP, CVN, RLK, MGA; Resources - AB, HRC, MAA; Data Curation - JM, JS, AB; Writing - Original Draft - AB, JS, and JM; Writing - Revision & Editing - All authors; Visualization - JM, JS, and AB; Supervision - AB and HRC; Project Administration - AB; Funding Acquisition - AB and HRC, additional fellowships in acknowledgments

## TABLE LIST

STable 1 - Primary Atlas Raw Metabolites

STable 2 - Primary Atlas Normalized Metabolites

STable 3 - Primary Atlas Isotopologues

STable 4 - Organoid Atlas Raw Metabolites

STable 5 - Organoid Atlas Normalized Metabolites

STable 6 - Organoid Atlas Isotopologues

STable 7 - Metadata for Glucose snRNA-seq

Stable 8 - Organoid Glucose Manipulation Atlas Raw Metabolites

STable 9 - Organoid Glucose Manipulation Atlas Normalized Metabolites

STable 10 - Organoid Glucose Manipulation Atlas Isotopologues

STable 11 - Metadata for PPPi snRNA-seq

Stable 12 - Organoid PPPi Atlas Raw Metabolites

STable 13 - Organoid PPPi Atlas Normalized Metabolites

STable 14 - Organoid PPPi Atlas Isotopologues

STable 15 - Metadata for shPPP scRNA-seq

Stable 16 - Organoid shPPP Atlas Raw Metabolites

STable 17 - Organoid shPPP Atlas Normalized Metabolites

STable 18 - Organoid shPPP Atlas Isotopologues

## METHODS

### Stem Cell Maintenance and Authentication

KOLF2 human induced pluripotent stem cell line (hIPSC) UCLA6 embryonic stem cell line (ESC)

UCLA1 embryonic stem cell line (ESC)

All stem cell lines were expanded on growth factor-reduced Matrigel-coated six-well plates. Stem cells were thawed and cultured with mTeSR Plus Basal Medium with mTeSR Plus 5x Supplement containing 1X penicillin/streptomycin. Medium was changed every day and lines were passaged when colonies reached 70% confluency. Lines were passaged with ReLeSR (CAT#100-0483) and cell lifters. Lines used for this study were between passages 15 - 30. Stem cells were karyotyped to ensure copy number integrity. Lines tested negative for mycoplasma prior to harvest.

### Generating cortical organoids

Cortical organoids are generated from an established protocol (Bhaduri et al., 2020; Pollen et al., 2019). Both PSC and ESC were expanded and dissociated with Accutase to a single-cell suspension. Cells were then reconstituted and aggregated with a neural induction medium containing Rock inhibitor Y-27632, SB431542 and IWR1-endo at a seeding density of 10,000 cells per well in a 96-well v-bottom low-attachment plate. Rock inhibitor is removed after 6 days. After 18 days, SB431542 and IWR1-endo are removed and organoids are transferred to a low-adhesion 6-well plate and an orbital shaker rotating at 100 rpm and changed to a medium containing DMEM/F-12 with GlutaMAX, 1x N2 supplement, CD Lipid Concentrate, and 1x penicillin-streptomycin. At day 35 the medium is changed to DMEM/F-12 with GlutaMAX, 1x N2 supplement, CD Lipid Concentrate, 1x penicillin-streptomycin, 10% fetal bovine serum (FBS), 5ug/mL Heparin, and 0.5% Matrigel. At day 70 the medium is DMEM/F-12 with GlutaMAX, 1x N2 supplement, CD Lipid Concentrate, 1x penicillin-streptomycin, 10% FBS, 5ug/mL Heparin, 0.5% Matrigel, and 1x B27 with vitamin A. Organoids were collected for Seahorse Analyzer, metabolomics and IF at week 5, 8, 10, 12, and 15.

### Human Primary Cortex Tissue (HPCT) Dissociation and Processing

HPCT is obtained and processed as approved by Ronald Reagan UCLA Medical Center Translational Pathology Core Laboratory. At the time of collection, HPCT is washed with 1X artificial cerebrospinal fluid (aCSF) containing NaCl, KCl, MgCl_2_-6H_2_O, CaCl_2_-2H_2_O, NaH_2_PO_4_- H_2_O, NaHCO_3_, and glucose. Cortex tissue is identified by anatomical and visual inspection and then isolated from the whole brain tissue. Cortex tissue is dissociated with Papain (Worthington)-containing DNase. Tissue with Papain and DNase mixture is incubated at 37 C incubator for 10 minutes and is followed by vigorous shaking every 5 minutes up until 40 minutes total. After tissue is broken up into small chunks, tissue is triturated to create a single-cell suspension. Cells are centrifuged at 300g for 5 minutes and resuspended with fresh media. An aliquot of final cell-suspension is used for cell counting.

Dissociated cells are plated at 1 million/mL in a 12-well poly-d-lysine (PDL) - coated tissue culture plate. For seahorse assays, cells were plated at 240,000/150uL on a 96-well Agilent Seahorse XFe96/XF Pro PDL Cell Culture Microplate (CAT#103799). Cells used for IF were plated at a 500,000/mL seeding density on a PDL Coated German Glass Cover Slip (CAT#72294). Dissociated HPCT cells were reconstituted in DMEM/F12-based medium containing 1x GlutaMAX, 1x B27, 1x N2, 1x sodium pyruvate and 1x penicillin-streptomycin. **I**mmunofluorescence (IF) and quantitative reverse transcription polymerase chain reaction (RT-qPCR) for cortex markers (FOXG1 and CTIP2) were used to confirm cortical identity.

Tissue collected was consented for research purposes by patients and samples were de-identified with no sex information known.

### RT-qPCR

RNA was isolated from HPCT and organoids using Qiagen RNeasy kits (CAT# 74104). Extracted RNA concentrations were measured using ThermoFisher NanoDrop and then converted into cDNA using a reverse transcriptase reaction from Invitrogen SuperScript™ IV VILO™ Master Mix (CAT#11756500) RT-qPCR. RT-qPCR reactions were performed in 384 well plates combining cDNA, gene-specific primers, and Power SYBR™ Green PCR Master Mix (CAT#4368708). Reactions were performed in triplicates for every gene probe. Samples were loaded and analyzed using a QuantStudio™ 5 Real-Time PCR System (CAT#A28140). Files were then exported and processed using Microsoft Excel. Signals for genes of interest were normalized to the signal of Glyceraldehyde-3-phosphate dehydrogenase (GAPDH) for each sample. Values were then converted into fold-change measurements of appropriate control groups for each experiment.

For detection of cortical signal from HPCT:

GAPDH

● Forward; 5’-tcaaggctgagaacgggaag-3’
● Reverse; 5’-cgccccacttgattttggag-3’ CTIP2
● Forward; 5’-tccagctacatttgcacaaca-3’
● Reverse; 5’-gctccaggtagatgcggaag-3’ FOXG1
● Forward; 5’-ccgcacccgtcaatgactt-3’
● Reverse; 5’-ccgtcgtaaaacttggcaaag-3’

For detection of PPP genes from genetic knockdown experiment G6PD

● Forward; 5’-tccagggcgatgccttccat-3’
● Reverse; 5’-gaaggccatcccggaacagc-3’ TKT
● Forward; 5’-tcaatcgcctgggccagagt-3’
● Reverse; 5’-ccacgctgtgtccatccacg-3’

### Methylation Data Collection and Analysis

Genomic DNA was harvested from week 8, 12 and 16 cortical organoids generated from multiple stem cell lines and HPCT samples ranging from gestational ages 14 - 34. The genomic DNA was processed for an Infinium MethylationEPIC Assay. The methylation data was then analyzed using the Fetal Brain Clock package on R (Steg et al., 2021) to estimate the methylation-based age of cortical organoids to HPCT. To convert post conceptional weeks to gestational weeks, two weeks were added to the original post conceptional week value.

### Seahorse Assay Analyzer

HPCT and cortical organoids were dissociated with Papain (Worthington)-containing DNase. Dissociated cells were plated at a 240k cells seeding density per well in an Agilent Seahorse XFe96/XF Pro PDL Cell Culture Microplate (CAT#103799). Seahorse plates were run with XFe96 Seahorse Extracellular Flux Analyzer in DMEM assay medium (Sigma D5030) supplemented with 5mM glucose, 2mM glutamine, 1mM pyruvate, and 5mM HEPES and included injection of compounds at final concentrations of 2uM oligomycin, 1.35uM FCCP, and 1uM rotenone, 2uM antimycin A, and 50mM 2-deoxy-D-glucose (2DG). The Seahorse Assay incorporates several compounds that inhibit or stimulate respiration and acidification rates that measure key metabolic responses. For oxygen consumption rates (OCR), the initial rate is known as the Basal OCR which is the endogenous rate of respiration. The first substrate introduced is oligomycin (Oligo), an ATP synthase inhibitor that reduces flow of electrons through the electron transport chain (ETC) in coupled mitochondria therefore causing a decrease in respiration. This decrease is known as ATP-Linked Respiration, the OCR linked to ATP production. The remaining rate during the Oligo treatment is known as Proton Leak, the rate independent of ATP synthase that is due to protons leaking across the inner membrane. Next, carbonyl cyanide-p-trifluoromethoxyphenylhydrazone (FCCP), an uncoupler, is introduced and activates the movement of protons across the inner membrane causing increased respiratory activity known as the Maximal Respiration. The difference rate between Maximal Respiration and Basal Respiration is known as the Reserve Capacity rate. Finally, Rotenone and Antimycin A (AA/Rot) are introduced as complex I and complex III inhibitors to fully block mitochondrial respiration displaying the remaining rate is known as the non-mitochondrial OCR. For the extracellular acidification rate (ECAR), the initial reading is known as the Basal ECAR which is the endogenous acidification rate. The introduction of Oligo, which inhibits ATP synthase and requires compensation in ATP synthesis through glycolysis, initiates an increase in ECAR and is correlated to the amount of lactate from glycolysis released in the extracellular medium when ATP synthase is inhibited. Finally, the introduction of 2-deoxy-D-glucose (2DG), a glucose analog, is a hexokinase inhibitor that shuts down glycolysis to provide the non-glycolytic acidification rate. Seahorse rates were normalized by cell count in each well, which is determined based upon Hoescht staining and imaging in an Operetta High Content Imaging System (PerkinElmer).

### Metabolomics Setup

Metabolite pool and metabolic pathway activity were analyzed via metabolomics and isotope ^13^C-tracing of both glucose and glutamine to acquire an extensive upstream and downstream metabolic overview. Gibco custom DMEM/F-12 (1:1) without D-glucose (dextrose), HEPES, L-glutamine, Phenol Red, Sodium Bicarbonate, and Sodium Pyruvate was supplemented with D-glucose (17.5mM) or D-glucose (U-^13^C_6_, 99%, CLM-1396), L-glutamine (2.5mM) or L-glutamine (^13^C_5_, 99%, CLM-1822), Sodium bicarbonate (2.438g/L) and Sodium Pyruvate (0.5mM). Isotopic ^13^C-tracing of D-glucose was supplemented into glucose-free custom DMEM/F-12 and Isotopic ^13^C-tracing of L-glutamine was supplemented into glutamine-free custom DMEM/F-12 supplement (glucose concentration changed for glucose manipulation experiment to 8.75mM or 2.5mM). Isotopic ^13^C-tracing of D-glucose and ^13^C-tracing of L-glutamine were administered to media of HPCT cells and cortical organoids 24 hours prior to metabolite extraction to ensure steady state of isotopic labeling. Samples were washed with 150mM ammonium acetate solution (pH 7.4) and metabolites were extracted with 80% methanol and 100nM trifluoromethanesulfonate (TFMS, internal standard). Samples were spun at 17,000 g (4 °C) for ten minutes to remove precipitated cell material (protein/DNA). Pellets containing DNA were stored in -20. Supernatants were collected, moved to a new clean tube, and then evaporated using a Nitrogen evaporator (Organomation). Metabolite extracts were processed and analyzed via Vanquish liquid chromatography Thermo Q-Exactive Orbitrap high mass accuracy (LCMS/MS) system to profile baseline metabolites in HPCT and the cortical organoid system.

Frozen pellets were resuspended in a solution containing 100mM NaCl, 20mM Tris-HCl (pH 7.4), 5mM EDTA, 0.1% SDS, Proteinase K (500 μg/mL). Resuspension volume was recorded to calculate total DNA in micrograms (ug). DNA concentration was measured using a nanodrop. Final concentration was determined by multiplying nanodrop concentration with resuspension volume.

### Metabolomics Analysis and Pathway Calculations

Metabolite measurements were performed as described in (Perez-Ramirez et al., 2024). Processed pool values were normalized by DNA concentration (ug) and TFMS values. An average of combined metabolite levels was calculated for each individual biological replicate after normalization. Each individual sample was then scaled to an arbitrary number to achieve an identical average combined metabolite level across each biological sample within a given dataset and experiment. Normalized and scaled values were used to perform comparisons and statistical tests. Isotopologues were normalized to Glucose M+6 values. Additionally, pathway calculations were also performed as described in interpretation of metabolite isotopologue distributions in Perez-Ramirez et al.

### Immunofluorescence (IF) protocol (for organoids and HPCT cells)

For immunofluorescence (IF) of cultured primary tissue samples, cells were fixed using 4% paraformaldehyde (PFA) for 15 minutes at room temp, then washed 3 times with PBS. Samples were then treated with a citrate-based antigen retrieval solution in PBS at 95°C for 20 minutes replacing PBS antigen retrieval solution 3 times. Samples were permeabilized and blocked in PBS supplemented with 0.1% Triton, 5% normal donkey serum, and 3% bovine serum albumin for 30 minutes at room temperature. Primary antibodies were diluted in antibody buffer (PBS supplemented with 0.1% Triton, 5% normal donkey serum, and 3% bovine serum albumin) and incubated with samples overnight at 4°C. Primary antibodies used include: CTIP2, SOX2, TBR2, GFAP, HOPX, SATB2, MAP2. Samples were washed 3 times using PBS with 0.1% Triton, then incubated with the appropriate secondary antibodies diluted in antibody buffer along with DAPI for 2 hours at room temperature away from light. After secondary incubation, samples were washed 3 times with PBS and mounted using Prolong gold antifade reagent.

For IF of cortical organoids, organoids were fixed using 4% PFA for 45 minutes at room temp, then incubated in 30% sucrose in PBS overnight. Organoids were then embedded into cryomolds using a 1:1 30% sucrose and OCT solution and stored at -80C. Cryomolds were sectioned using a cryostat at 16um slices. Samples were then treated with a citrate-based antigen retrieval solution in PBS at 95°C for 20 minutes replacing PBS antigen retrieval solution 3 times. Samples were permeabilized and blocked in PBS supplemented with 0.1% Triton, 5% normal donkey serum, and 3% bovine serum albumin for 30 minutes at room temperature. Primary antibodies were diluted in antibody buffer (PBS supplemented with 0.1% Triton, 5% normal donkey serum, and 3% bovine serum albumin) and incubated with samples overnight at 4°C. Primary antibodies used include: CTIP2, SOX2, TBR2, GFAP, HOPX, SATB2, MAP2. Samples were washed 3 times using PBS with 0.1% Triton, then incubated with the appropriate secondary antibodies diluted in antibody buffer along with DAPI for 2 hours at room temperature away from light. After secondary incubation, samples were washed 3 times with PBS, and coverslips were mounted using Prolong gold antifade reagent.

SOX2

● Santa Cruz
● SC-365823
● Mouse
● 1:500

CTIP2

● Abcam
● AB18465
● Rat
● 1:500

GFAP

● Abcam
● AB4674
● Chicken
● 1:500

TBR2

● R&D systems
● AF6166
● Sheep
● 1:500

HOPX

● Proteintech
● 11419-1-AP
● Rabbit
● 1:500

MAP2

● LSBIO
● C61805
● Chicken
● 1:500

SATB2

● Abcam
● AB51502
● Mouse
● 1:500

GFP

● Abcam
● AB4674
● Chicken
● 1:500

KI67

● Abcam
● AB16667
● Rabbit
● 1:500

Cleaved-caspase 3

● Cell Signaling Technology
● 9664S
● Rabbit
● 1:500

PAX6

● Abcam
● AB78545
● Mouse
● 1:500

### EVOS Imaging and Imaris Quantification (for organoids and HPCT cells)

Coverslips were imaged on EVOS 5000 (Thermo) fluorescent microscope using appropriate excitation/emission lasers and filters for the corresponding immunofluorescent antibodies and were matched across comparable conditions. Individual channels were saved as Tiff files and merged together into a composite image on FIJI. The merged channel image file was then converted into an Imaris file using ImarisFileConverter. Merged channel Imaris images were quantified using the “spots” function to identify DAPI positive cells and then classified as positive for other markers based on signal intensity of other channels. A positive signal intensity for a gene was determined by thresholding the mean intensity or sum of square of a channel within each spot.

### Glucose Experiment Setup for Organoids

Cortical organoids are generated as previously described in the “Generating cortical organoids” section. Gibco custom DMEM/F-12 (1:1) without D-glucose (dextrose), HEPES, L-glutamine, Phenol Red, Sodium Bicarbonate, and Sodium Pyruvate was supplemented with D-glucose (17.5mM, 8.75mM or 2.5mM), Sodium bicarbonate (2.438g/L) and Sodium Pyruvate (0.5mM). At week 5, cortical organoids are cultured in a custom DMEM/F-12-based medium supplemented with 2.5mM GlutaMAX, 10% FBS, 5ug/mL Heparin, 1x N2, and 1x CD Lipid Concentrate and 0.5% Matrigel. Cortical organoids are divided into four condition groups: control baseline glucose concentration (17.5mM), supplemented 17.5mM glucose, supplemented 8.75mM glucose, and supplemented 2.5mM glucose. At day 70, organoids are moved to a medium supplemented with 2.5mM GlutaMAX, 1x N2 supplement, CD Lipid Concentrate, 1x penicillin-streptomycin, 10% FBS, 5ug/mL Heparin, and 0.5% Matrigel and previously described glucose condition groups. Cortical organoids are harvested at week 12 and processed for Seahorse Analyzer, metabolomics, IF and snRNA-seq. Throughout organoid culture, media was replenished every 2 days.

### Pentose Phosphate Pathway pharmacological inhibitors for Organoids

Cortical organoids are generated as previously described in “Generating cortical organoids” section. At week 8, cortical organoids are cultured in a DMEM/F-12-based medium containing 10% FBS, 5ug/mL Heparin, 1x N2, and 1x CD Lipid Concentrate and 0.5% Matrigel. Cortical organoids are divided into four condition groups: control baseline, DMSO, 1uM 6- Aminonicotinamide (6AN, CAT# HYW010342), and 37.5uM Oroxylin A (CAT#HY-N0560). All inhibitors will be reconstituted in DMSO and will be added to the media of cultured cortical organoids at an equivalent 0.05% concentration across all conditions with a matched volume vehicle control of DMSO and untreated organoids as a control. At day 70, organoids are moved to a medium supplemented with 1x N2 supplement, CD Lipid Concentrate, 1x penicillin-streptomycin, 10% FBS, 5ug/mL Heparin, and 0.5% Matrigel and previously described with previously described PPP-inhibitor condition groups. Cortical organoids are harvested at week 12 and processed for metabolomics, IF, and snRNA-seq. Throughout organoid culture, media was replenished every 2 days.

### Generation of a SOX2 enhancer-driven EGFP reporter

We surveyed the literature for regulatory elements that can be used to drive EGFP expression in a SOX2-dependent manner in cortical radial glia and identified SRR2, a 200-bp SOX2 enhancer located approximately ∼ 4-kb downstream of the SOX2 transcription start site. SRR2 is active in embryonic and telencephalic neural stem cells but undergoes methylation-dependent silencing as NSCs differentiate into postmitotic neurons (Miyagi et al., 2006; Miyagi et al., 2004; Sikorska et al., 2008). Benchmarking against two independent single-cell/nuclei epigenomics datasets profiling the developing human cortex confirmed that SRR2 is selectively accessible in radial glia cells (Wang et al., 2025; Ziffra et al., 2021). To generate an SRR2-driven EGFP reporter, SRR2 was first PCR-amplified from human genomic DNA using primers with 20-bp homology arms for subsequent plasmid assembly (forward primer: 5’- cactttggcgccggcttcccccctaattaatgcagaga-3’; reverse primer: 5’- ccattatataccctctactagcccctgcctcaaattccgggaat-3’). Next, the existing hPGK promoter in a Gateway-compatible pLEX-305 lentiviral plasmid (Addgene 41390) was excised using XhoI, and the SRR2 PCR product was cloned into the linearized backbone using the NEBuilder HiFi DNA Assembly reaction (New England Biolabs E2621L). This yielded the pLEX-305-SRR2-minP- GW-WPRE destination vector, which contains an attR1/2-flanked Gateway cloning site (GW) downstream of the SRR2 enhancer coupled to a minimal promoter (minP) sequence (5’- tagagggtatataatggaagctcgacttccag-3’) followed by the woodchuck hepatitis virus posttranscriptional regulatory element (WPRE) at the 3’ UTR for enhanced transgene expression. An EGFP cassette was then subsequently cloned from a pDONR221-EGFP entry clone into the Gateway site using a standard LR reaction.

### Isolation of Radial Glia Using SRR2-SOX2 enhancer

SRR2/EGFP plasmids were packaged into lentivirus and infected into all 3 stem cell lines before aggregation into cortical organoids. SRR2/EGFP organoids were cultured as previously mentioned above. Starting at week 8 organoids were dissociated as discussed previously and prepped for FACS. Prior to sorting on BioRad S3e cell sorter, organoids were resuspended in FACS buffer (HBSS with 1% BSA, 0.1% glucose, and 0.2 mM EDTA) with DAPI. Cells and debris removal were selected based on forward and side scatter area, followed by two-step doublet removal using forward and side scatter height and area. Live cells were selected based on negative signal from DAPI and SRR2+ radial glia cells were selected based on log fold change of signal intensity from EGFP. Cells were sorted into appropriate FACS tubes and then resuspended in media used for 2D culturing of HPCT and subsequently plated onto PDL-coated plates.

### Intracellular staining of SRR2/EGFP+ organoids

SRR2/EGFP organoids were generated as described above. Starting at week 8 organoids were dissociated using Papain (Worthington)-containing DNase. Cells were resuspended in FACS buffer (HBSS with 1% BSA, 0.1% glucose, and 0.2 mM EDTA). Organoid cells were incubated with Zombie Violet from BioLegend Fixable Viability Kit (CAT#423114) and subsequently fixed in 4% PFA. Cells were then permeabilized using 0.1% Triton and stained using primary antibodies conjugated to fluorophores targeting either SOX2 or CTIP2. Cells were immediately analyzed using Invitrogen Attune NxT Flow Cytometer. Analysis of flow cytometry files were performed on FlowJo. Selection and debris removal were based on forward and side scatter area, followed by two-step doublet removal using forward and side scatter height and area. Live cells were selected based on a negative signal from Zombie Violet. SRR2/EGFP, SOX2, and CTIP2 cells were chosen based on log fold change of signal intensity from EGFP or secondary antibodies.

### Pentose Phosphate Pathway Knockdown for Organoids

To knockdown PPP genes, the pLKO.1 plasmid was used to create the shRNA targeting TKT or G6PD and obtained from Addgene (Plasmid# 10878). The puroR sequence was removed and replaced with mCherry via NEBuilder HiFi DNA Assembly reaction (New England Biolabs E2621L). The scramble (5’-cctaaggttaagtcgccctcg-3’), TKT (5’-cgccgaactgctgaagaaagaa-3’), and G6PD (5’-gcctcagtgccacttgaca-3’) sequences were inserted into the backbone according to manufacturer’s instructions. TKT and G6PD sequences were obtained from published studies (Li et al., 2009; Qin et al., 2019).

To produce lentivirus from these plasmids, HEK293T cells were transfected with each plasmid alongside both psPAX2 and pMD2.G using Lipofectamine 2000 (Invitrogen 11668019). Media containing the lentivirus was concentrated using Lenti-X Concentrator (Takara Bio 631232).

Control organoids were dissociated and plated onto coated PDL plates as discussed above. Plated organoid cells were infected with lentivirus containing shRNA targeting either TKT, G6PD or appropriate scramble control. RNA knockdown of this system was validated by RT-qPCR and relative RNA fold change expression was compared to scramble for each experiment. For radial glia isolation and PPP knockdown SRR2/EGFP organoids were dissociated, underwent FACS for EGFP+ cells, and plated with lentivirus for production of shTKT or shG6PD. Metabolic and scRNA-seq analysis was performed after 7 days in culture post infection.

### Statistical Analysis

Significance between groups for immunofluorescence quantification, seahorse analysis, metabolite abundance and labeling were performed in Graphpad Prism using Brown-Forsythe and Welch ANOVA tests and unpaired t test with Welch’s correction and with individual variances computed for each comparison. Significance was indicates by the following asterisks: * indicates p-value < 0.05, ** indicates p-value < 0.01, *** indicates p-value < 0.001, **** indicates p-value < 0.0001.

P-value significance for differential analysis of LCMS-based metabolomics in HPCT and organoid was determined by performing calculations in excel between groups selected for comparison. Calculations for p-value were computed by performing -log_10_ of the t test between all biological and technical replicates in a specific DP compared to another DP. A two-tailed t test with two-sample unequal variance was used.

Statistical analysis for module scores was performed using a t-test in R to compare distributions between large sample sizes that fall into a classic normal distribution.

### Single-nuclei and single-cell RNA-sequencing capture and analysis

Single-nuclei capture from frozen cells for the organoid manipulation and PPPi experiments was performed following Chromium’s Nuclei Isolation Kit followed by the Chromium GEM-X v4 protocol. Single-cell capture from live cells for the shRNA targeting PPP enzyme experiment was performed following Chromium GEM-X v4 manufacturer’s instructions for organoids. Batch is indicated in the metadata annotation in Supplemental Table 7, 11, and 15. In the single-nuclei experiments, 20,000 cells were targeted for capture and 12 cycles of amplification for each of the cDNA amplification and library amplification were performed. For a single-cell experiment for shPPP, 10,000-20,000 cells were targeted for capture and 12-13 cycles of amplification for each of the cDNA amplification and library amplification were performed. Libraries were sequenced as per manufacturer recommendation on a NovaSeq X flow cell.

Fastq files were aligned using cellranger v8 and were processed with CellBender to remove ambient RNA (Fleming et al., 2023). DoubletFinder (McGinnis et al., 2019) was used to remove doublets from the data. Data was then analyzed using Seurat v5 (Hao et al., 2024). Quality control removed cells with fewer than 500 genes per cell and more than 10% mitochondrial content. Clustering was performed using standard Seurat workflows. Cell types were defined by taking the highest predicted cell id from Seurat’s MapQuery analyses with a recent developmental meta-atlas as the reference (Hao et al., 2024). Cell types were aligned to “Subtype” and then were simplified based upon markers of upper versus deep layer neurons in the feature plots presented in supplemental figures. The fraction of these simplified cell types was used to calculate fold changes of cell types. Differential gene expression across conditions was performed with Seurat using a Wilcoxon rank-sum test and were analyzed for interesting ontology programs using Enrichr (Kuleshov et al., 2016).

Module analysis of KEGG pathways and from published data was performed using the ModuleScore feature from Seurat and plotted using ggplot2 in R.

---

**Supplemental Figure 1.**
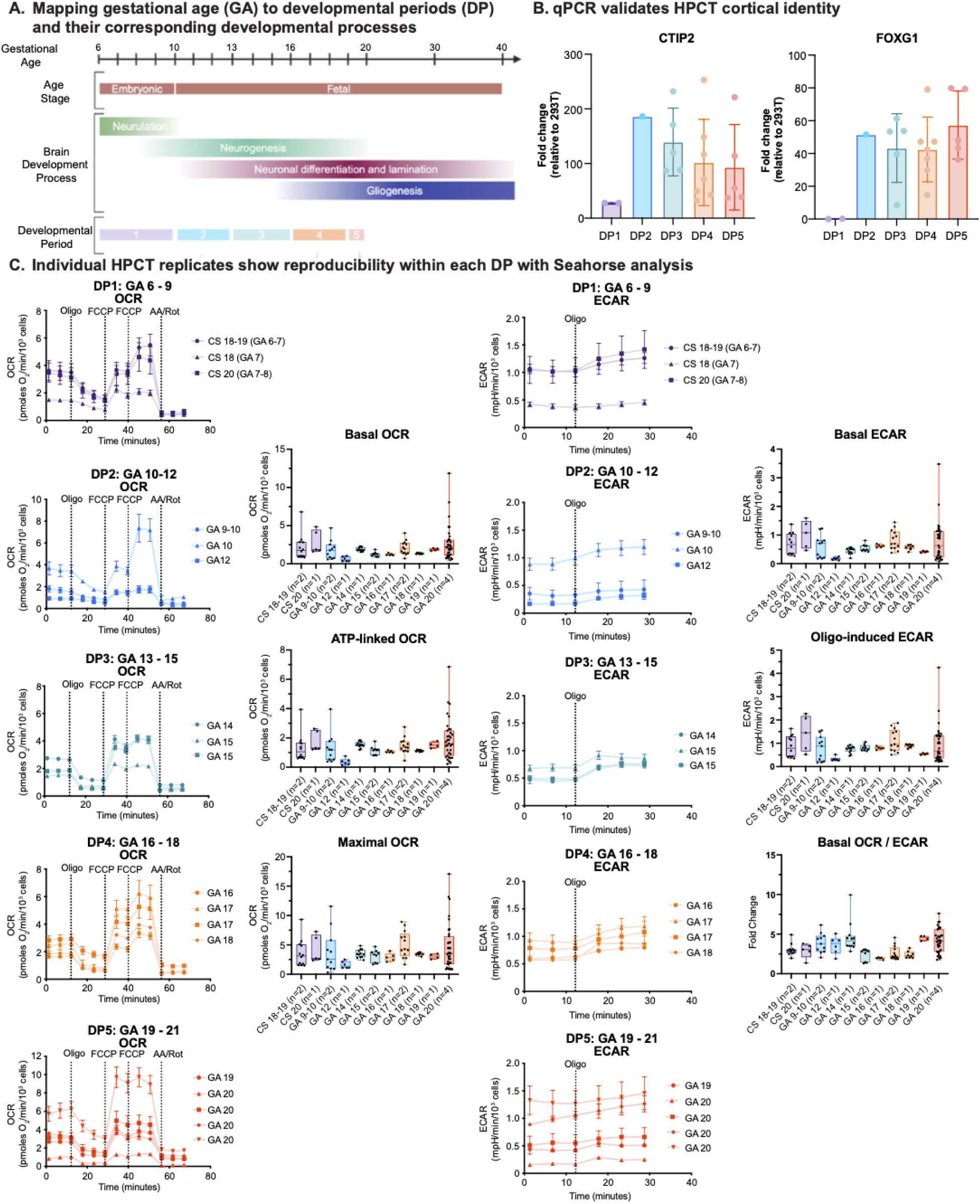
Human Primary Cortical Tissue Processing Pipeline Captures Expected Cortical Signatures and Allows for Confident Metabolic Analysis Across DPs. A. Schematic of developmental periods (DP) of interest and their corresponding gestational age (GA) as well as their corresponding brain developmental processes. During DP1, consisting of GA 6-9, neurulation occurs, and neurogenesis begins. In DP2, consisting of GA 10-12, neurogenesis increases while neuronal differentiation and lamination begins. In DP3, consisting of GA 13-15, neurogenesis, neuronal differentiation and lamination continues while gliogenesis begins. In DP4, consisting of GA 16-18, and DP5, and GA 19-20, neurogenesis, neuronal differentiation, lamination, and gliogenesis continue to increase. DP1 is considered an embryonic stage while DP2 onwards are considered fetal stages. B. To validate that our HPCT samples are cortex specific, we collected RNA from tissue samples incorporated in our analysis and quantified cortex-specific gene expression through quantitative reverse transcription polymerase chain reaction (RT-qPCR). Genes used to validate cortex tissue include CTIP2 and FOXG1. Gene expression quantification is displayed as the fold change in respect to negative control HEK293T cells. Error bars represent standard error of mean (SEM). C. Seahorse analysis was performed to measure oxygen consumption rate (OCR) and to calculate OCR response rates (left two columns, line graphs and box and whisker plots) as well as extracellular acidification rate (ECAR) and acidification response rates (right two columns, line graphs and box and whisker plots) for each HPCT sample within each DP. Basal OCR measures the endogenous respiration rate, ATP-linked OCR measures the respiration associated with ATP production after the addition of Oligo, and Maximal OCR measures the maximal respiration rate as a response to the protonophore FCCP. Basal ECAR measures endogenous acidification rate and is a proxy for glycolytic activity, Oligo-induced ECAR measures the rise in acidification in response to Oligo being introduced, and the basal OCR/ECAR rate is a calculation that measures the fold change amount of basal OCR activity compared to basal ECAR activity. Both OCR and ECAR are normalized to the number of cells in the plate, which is determined based upon Hoescht staining high-content imaging. In DP1, three samples consisting of the age: GA6-7, GA7, and GA7-8 were included. In DP2, three samples consisting of the ages: GA9-10, GA10, and GA12 were included. In DP3, three samples consisting of the ages: GA14, GA15 and GA15 were included. In DP4, four samples constant of the ages: GA16, GA17, GA17, and GA18 were included. In DP5, five samples consisting of the ages, GA19 and four GA20 samples were included. Each sample within each DP consisted of 6 - 12 technical replicates. Error bars represent SEM. Significance was calculated on the combined dataset presented in Figure 1.

**Supplemental Figure 2:**
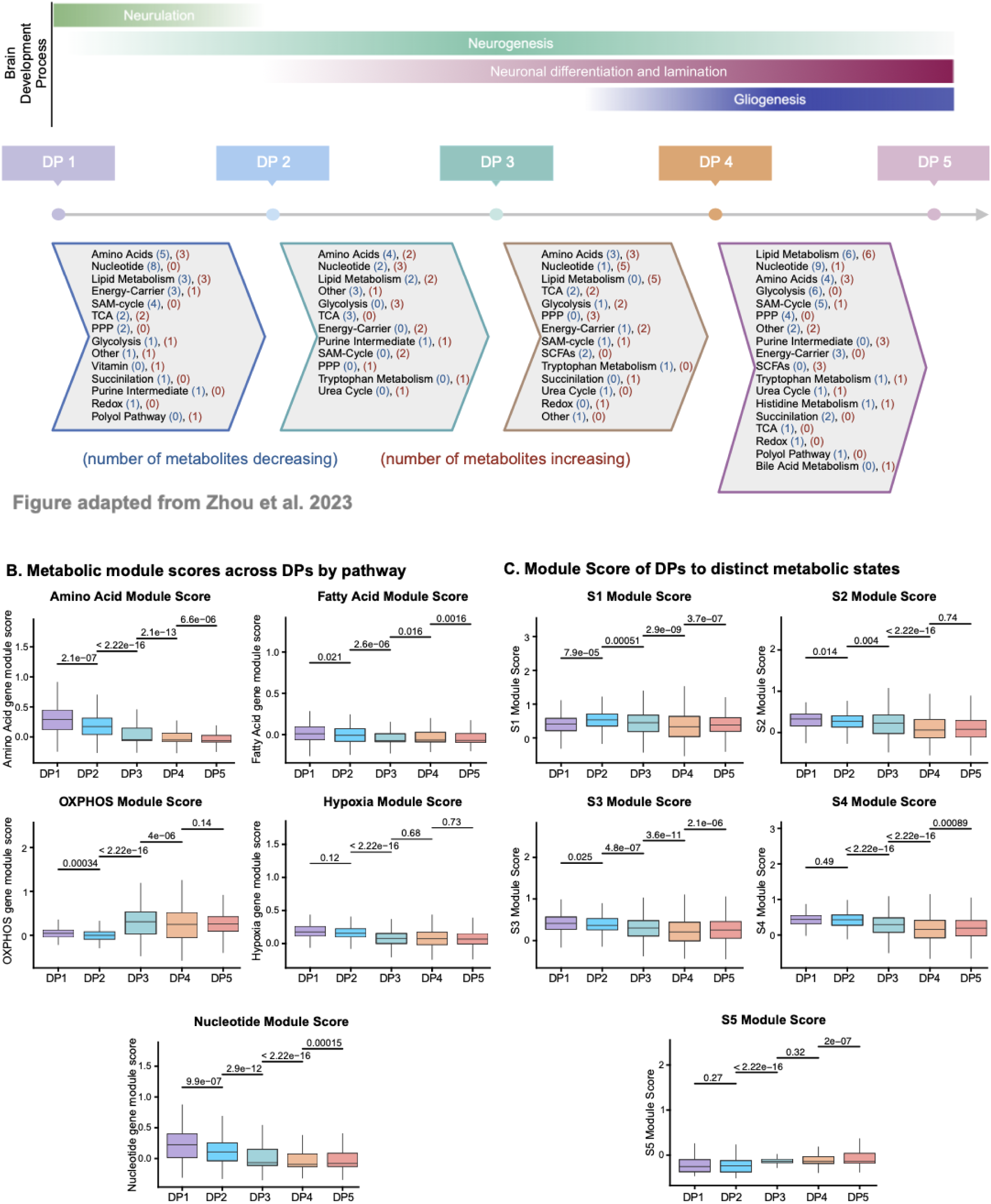
Investigating Metabolite Dynamics Across Human Cortical Developmental Periods. A) Schematic of DP timeline alongside brain developmental processes known to occur throughout DP1 - DP5. Chevrons indicate differential metabolites organized by pathways determined by log_2_ fold change from previous DP to subsequent DP. P-values are determined by t-test analysis. Each DP consists of three to five biological replicates with three to six technical replicates per sample. Pathways are followed with blue numbers that indicate the number of metabolites that were significantly decreased in DPx vs DPx- 1. Red numbers indicate the number of metabolites that were significantly increased in each comparison. B) Modules of metabolism genes were examined across developmental periods by leveraging a module score (Hao et al., 2024) to look at gene activity over time. Gene lists were obtained from KEGG pathway analysis and show significant dynamic changes over time, some of which match metabolic changes in Fig 2 but also many differences are also observed. C) Module scores from a recently published mouse analysis of metabolism genes during mouse development (Dong et al., 2022). These scores show dynamic gene programs that are similar to what was observed in mouse and correspond to some of the same metabolomic dynamics in Fig 2.

**Supplemental Figure 3:**
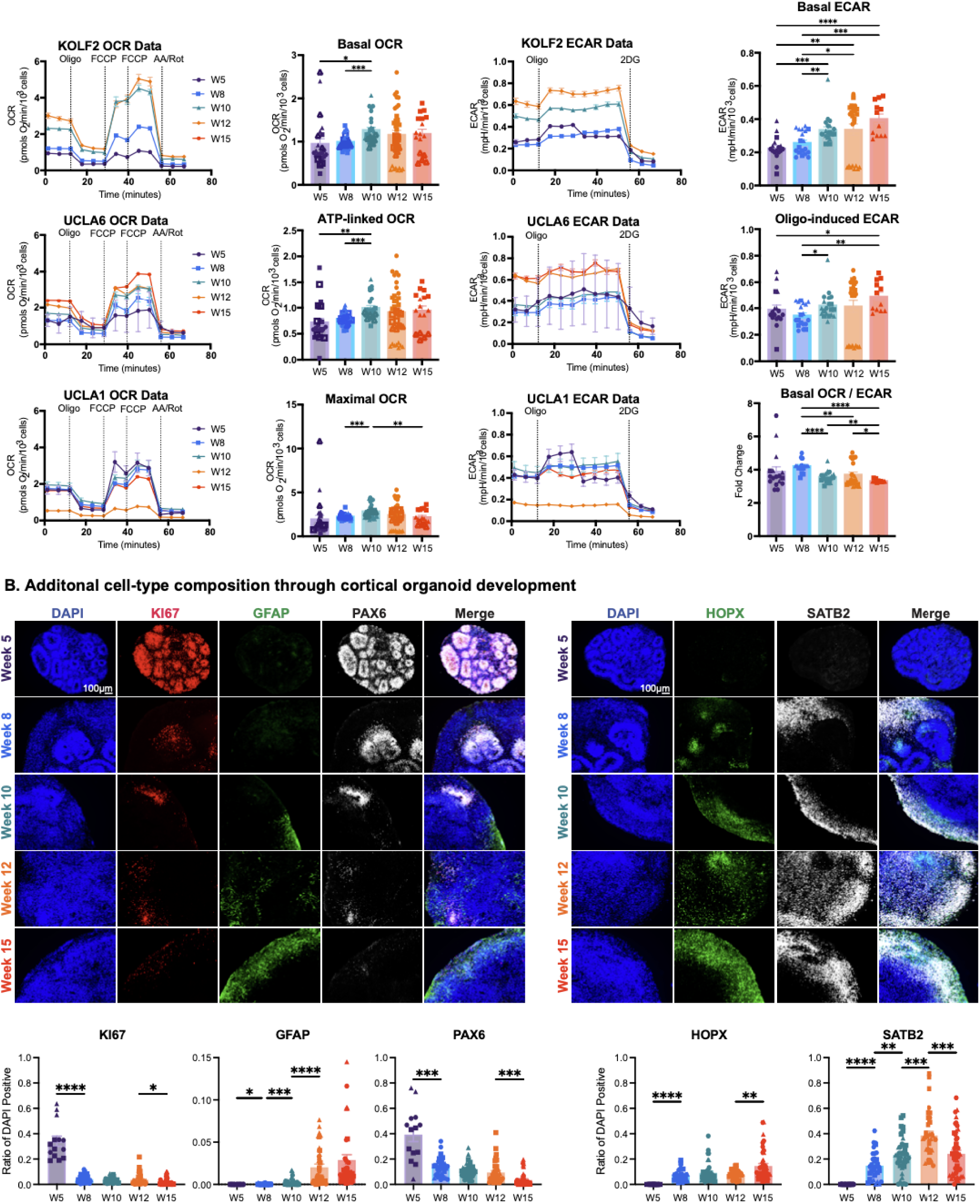
Metabolic Patterns in Cortical Organoids are Similar to HPCT Metabolic Trends. A) Seahorse analysis was performed to measure oxygen consumption rate (OCR) and to calculate OCR response rates (left two columns, line graphs and bar graphs) as well as extracellular acidification rate (ECAR) and acidification response rates (right two columns, line graphs and bar graphs) across three PSC lines (KOLF2, UCLA6, and UCLA1) with 5 time points- Week 5 (W5), week 8 (W8), week 10 (W10), week 12 (W12), and week 15 (W15). Basal OCR measures the endogenous respiration rate, ATP-linked OCR measures the respiration associated with ATP production after the addition of Oligo, and Maximal OCR measures the maximal respiration rate as a response to the protonophore FCCP. Basal ECAR measures endogenous acidification rate and is a proxy for glycolytic activity, Oligo-induced ECAR measures the rise in acidification in response to Oligo being introduced, and the basal OCR/ECAR rate is a calculation that measures the fold change amount of basal OCR activity compared to basal ECAR activity. Both OCR and ECAR are normalized to the number of cells in the plate, which is determined based upon Hoescht staining and high-content imaging. Three organoid replicates of each time point per line were analyzed, with 6-12 technical replicates for each time point. Error bars represent SEM. Significance was calculated with an ANOVA and a Tukey’s multiple comparisons test. ** indicates p-value < 0.01, **** indicates p- value < 0.0001. B) To validate that cell-type composition mirrored expected patterns as in HPCT, we performed immunofluorescence on cortical organoids. A representative image from the KOLF2 line for each of the 5 time points is shown in both panels here. Left panel shows staining for KI67 (red), a division marker, staining for GFAP (green) astrocyte populations, and staining for PAX6 (white) stains for neural progenitor cells (NPCs) and radial glia. Right panel shows staining for HOPX, a proxy for outer radial glia (oRG) and staining for SATB2 (white) marking upper-layer neurons. DAPI, shown in blue, marked nuclei. Scale bar is 100 μM and reflects all images. Quantification of the ratio of DAPI positive cells marked by each protein is presented in bar charts to the right, where three distinct biological replicates were analyzed for time point, and three technical replicates were used within each biological replicate; shapes on the bar chart correspond to each of the cell lines used (KOLF2 - circles, UCLA6 - square, UCLA1 - triangle). Significance was calculated using the Brown-Forsythe and Welch ANOVA. * indicates p-value < 0.05, ** indicates p-value < 0.01, *** indicates p-value < 0.001.

**Supplemental Figure 4:**
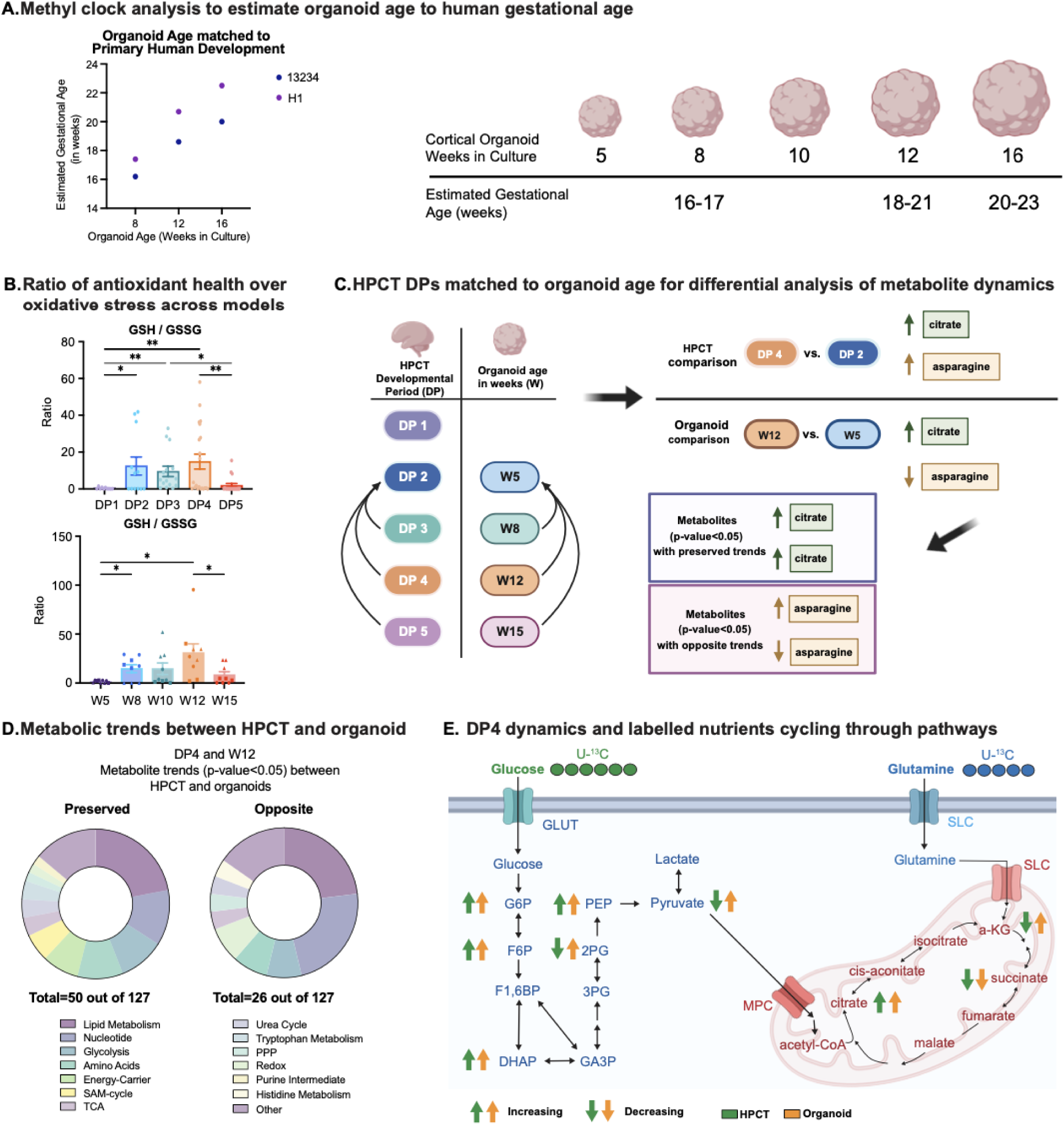
Aligning Metabolic Cortical Organoid Development to HPCT. A) Methyl clock analysis of cortical organoids harvested at weeks 8, 10, and 16 in culture. One biological replicate of each week was analyzed with one technical replicate in H1 (hESC) and 13234 (iPSC) stem cell lines. Methylation data was processed using a previously described methyl clock pipeline to determine the estimated gestational age of cortical organoids corresponding to HPCT development (Steg et al., 2021). Schematic showing organoids harvested at weeks 8, 10, and 16 were calculated to have an estimated gestational age of 14-15, 16-19, and 18-21, respectively. B) Bar graphs of the ratio of antioxidant health over oxidative stress across HPCT and organoid DPs. Three to five biological replicates of each DP were analyzed, with three to six technical replicates for each sample. Y-axis indicates average metabolite abundance of glutathione (GSH) over glutathione disulfide (GSSG). Dots show technical replicates from each biological sample within DPs. (KOLF2 - circles, UCLA6 - square, UCLA1 - triangle) Significance was calculated using the Brown-Forsythe and Welch ANOVA. * indicates p-value < 0.05, ** indicates p-value < 0.01, *** indicates p-value < 0.001. Error bars represent SEM. C) Schematic aligning organoid age in weeks (W) to DPs in HPCT for appropriate metabolomic comparison between systems. Week 5, 8, 12, and 15 organoids correspond to DPs 2, 3, 4, and 5, respectively. Alignment of DP1 to organoid age was not captured in our experimental setup. Matching DPs allows for differential analysis of metabolites at specific time points of neuronal development to determine significantly preserved or opposing trends between systems. Metabolite abundance for each metabolite was calculated by averaging all technical replicates across biological samples within DP2 and DP4. For instance, citrate is increasing in both HPCT and organoid at DP4 compared to DP2, while asparagine is increasing in HPCT but decreasing in organoid. Significantly trending metabolites across development were determined by p-value lower than 0.05. D) Circular graphs of preserved and opposite metabolic trends between HPCT and organoid at DP4 organized by metabolic pathway. Three to five biological replicates of each DP were analyzed, with three to six technical replicates for each sample. 50 out of 127 targeted metabolites dynamics were preserved between HPCT and organoid at DP4 while 26 metabolites exhibited opposite dynamics. Significantly changing metabolites were determined by p-value lower than 0.05. E) Schematic of ^13^C_6_-glucose and ^13^C_5_-glutamine cycling through glycolysis and the TCA cycle. Downstream ^13^C-label derived from ^13^C_6_-glucose label is highly abundant in glycolysis while downstream ^13^C-label derived from ^13^C_5_-glutamine label is primarily abundant in the TCA cycle. Upward and downward arrows represent metabolites whose abundance at DP4 vs DP2 and W12 vs W5 in HPCT and organoid increases or decreases, respectively. Green and orange arrows indicate metabolite dynamics in HPCT and organoid, respectively.

**Supplemental Figure 5:**
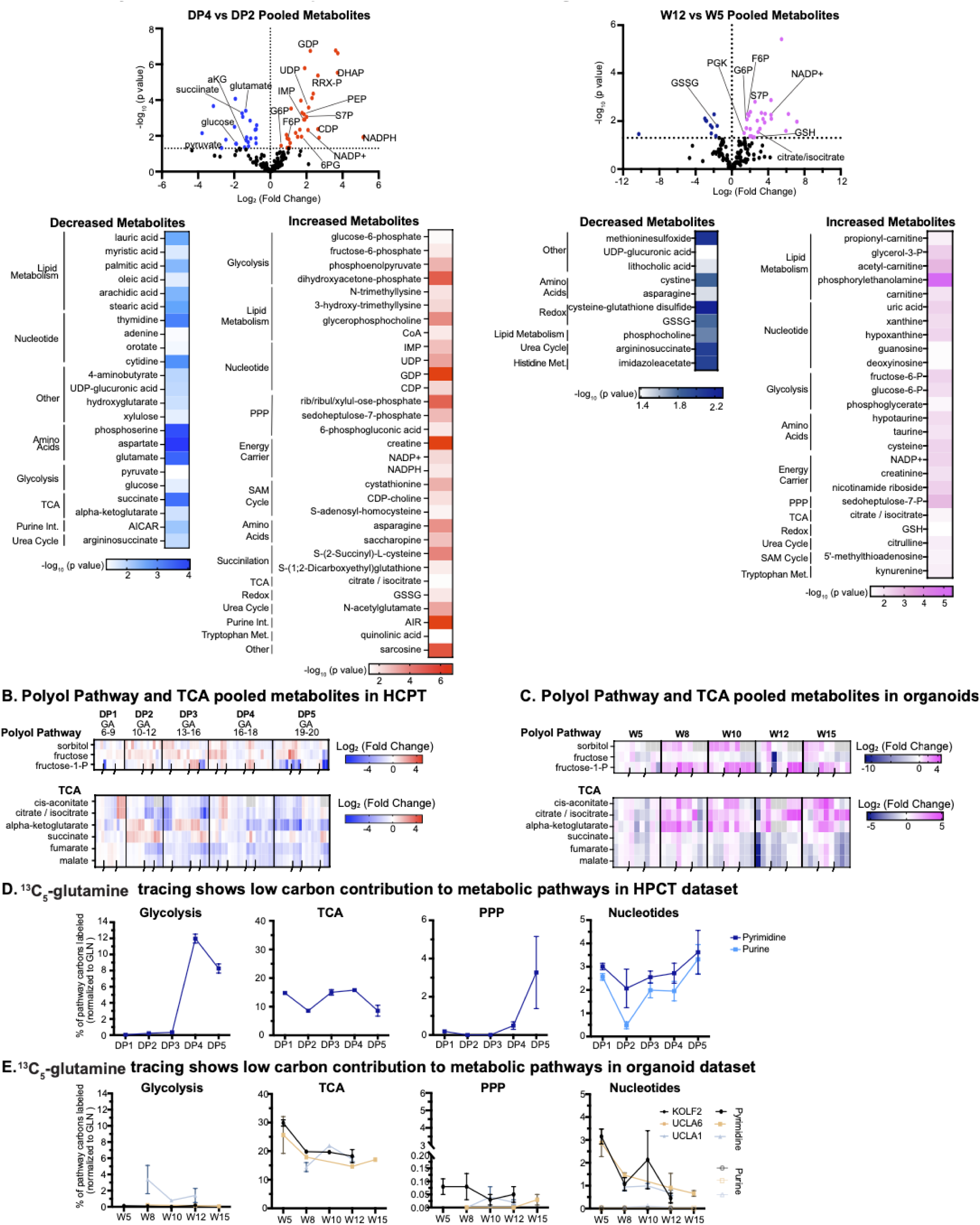
HPCT and Cortical Organoids Metabolomics Show Similar Trends. A) Volcano plots and heatmaps of differential metabolite abundance between DP4 vs DP2 in HPCT (left volcano plot, blue and red heatmaps) and W12 vs W5 in organoid (right volcano plot, dark blue and magenta heatmaps) show dynamic changes in pooled metabolomics across cortical development. The x-axis measures log_2_ fold change of pooled metabolite differences between the two time points within HPCT and organoids. The Y-axis measures significance of differential metabolite abundance represented by - log_10_ of p-value. Heatmaps list the differentially abundant metabolites between time points of interest and are organized by pathway. Heatmap values show the level of significance represented by -log_10_ of p-value. Red and pink heatmaps indicate significantly upregulated metabolites, and blue and dark blue vindicate significantly downregulated metabolites, in HPCT and organoids, respectively. Three to five biological replicates of each time point were analyzed within each system with three to six technical replicates for each biological replicate. P-values are determined by t-test analysis. B) Heatmaps of additional pathways that branch from glycolysis: the polyol pathway (top) and the tricarboxylic acid (TCA) cycle (bottom). Heatmaps display metabolite abundance levels from DP1 to DP5 and are determined by a log_2_ fold change to GA6. Three to five biological replicates of each DP were analyzed, with three to six technical replicates for each sample. C) Heatmap of polyol-pathway (top) and TCA (bottom) intermediates in cortical organoids. Values reflect abundance levels from W5 - W15 and were calculated by log_2_ fold change to W5. Three cell lines, with three organoids per cell line, for each time point were analyzed. D) HPCT pathway analysis of ^13^C_5_-glutamine label routed to glycolysis, PPP, and nucleotide metabolism. Three to five biological replicates of each DP were analyzed, with three to six technical replicates for each sample. Metabolites in glycolysis analysis include G6P, F6P, F16BP, DHAP, 3PG, PEP, and Pyruvate. Metabolites in TCA analysis include cis-aconitate, citrate/isocitrate, alpha-ketoglutarate, succinate, fumarate, and malate. Metabolites in PPP include 6PG, RRX5P, and S7P. Metabolites in purine metabolism (light blue) include hypoxanthine, xanthine, uric acid, AICAR, IMP, inosine, AMP, ADP, ATP, adenine, adenosine, GMP, GDP, GTP, guanine, and guanosine. Metabolites in pyrimidine metabolism (dark blue) include carbamoyl-aspartate, dihydroorotate, orotate, UMP, UDP, UTP, CMP, CDP, CTP, cytidine, and uridine. Y-axis measures the percent of total carbons in pathways that contained ^13^C-label normalized to ^13^C-glucose label. Error bars represent SEM. E) Organoid pathway analysis of ^13^C_5_-glutamine label routed to glycolysis, PPP, and nucleotide metabolism. Three cell lines were used, with 3 organoid per cell line for each time point. Metabolites in glycolysis analysis include G6P, F6P, F16BP, DHAP, 3PG, PEP, and Pyruvate. Metabolites in TCA analysis include cis-aconitate, citrate/isocitrate, alpha-ketoglutarate, succinate, fumarate, and malate. Metabolites in PPP include 6PG, RRX5P, and S7P. Metabolites in purine metabolism include hypoxanthine, xanthine, uric acid, AICAR, IMP, inosine, AMP, ADP, ATP, adenine, adenosine, GMP, GDP, GTP, guanine, and guanosine. Metabolites in pyrimidine metabolism include carbamoyl-aspartate, dihydroorotate, orotate, UMP, UDP, UTP, CMP, CDP, CTP, cytidine, and uridine. Y-axis measures the percent of total carbons in pathways that contained ^13^C- label normalized to ^13^C_5_-glutamine label. Pyrimidines and purines in nucleotide metabolism are separated by dark and light colors, respectively (KOLF2 - black circles, UCLA6 - orange squares, UCLA1 - grey triangle). Error bars represent SEM.

**Supplemental Figure 6:**
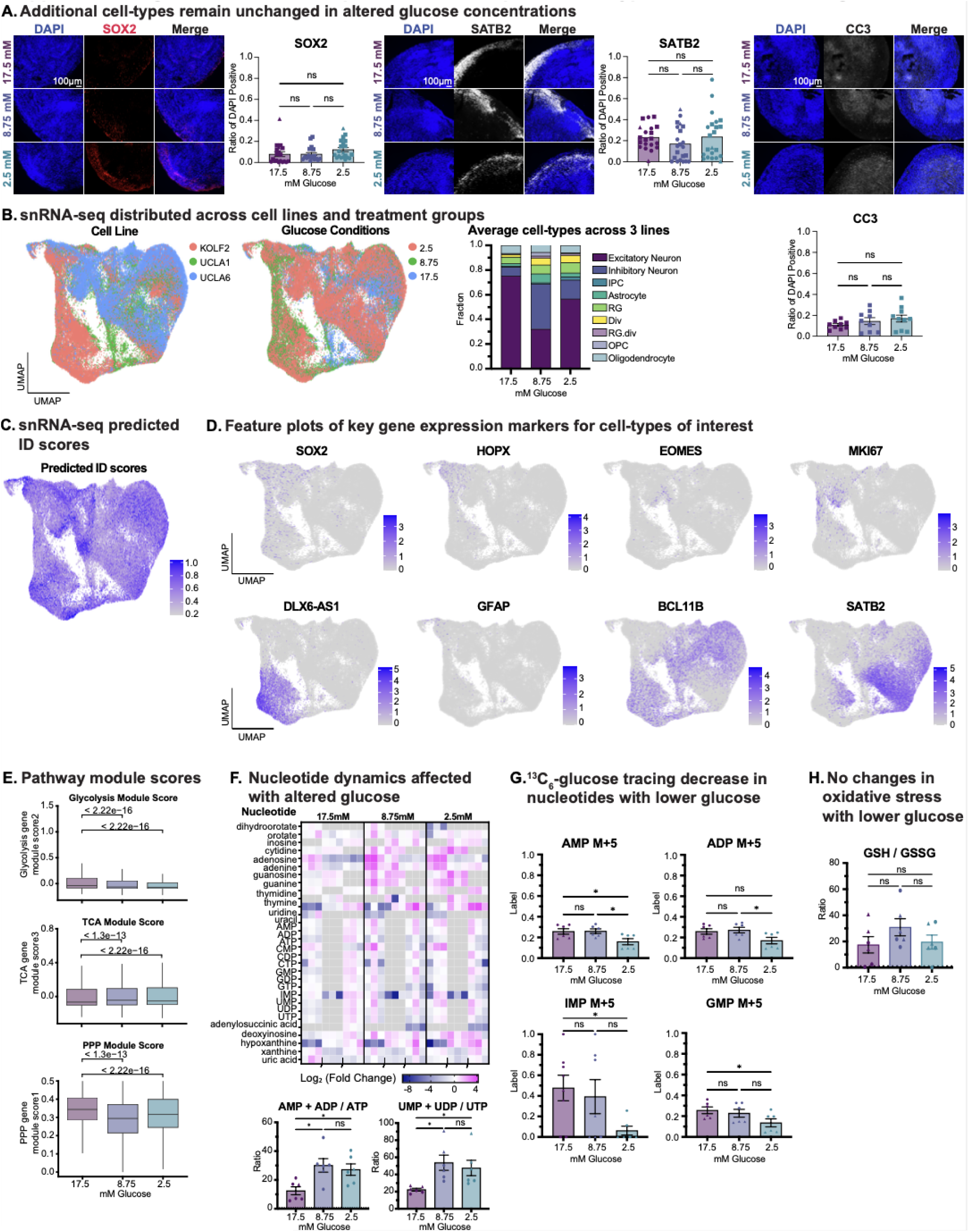
Glucose Manipulation Exhibit Specific Cell Type and Metabolic Changes. A) Immunofluorescence images of SOX2, SATB2 and CC3 in glucose manipulated organoids. A representative image from the KOLF2 line is shown for SOX2 and SATB2 and CC3 is represented by the UCLA6 line. Staining for CC3 labels apoptotic cells and negative signal is indicative of positive cell health. DAPI, shown in blue, marked nuclei. Scale bar is 100 μM and reflects all images. Quantification of the ratio of DAPI positive cells marked by each protein is presented in bar charts to the right, where up to three cell lines with two to three distinct organoid replicates per line were analyzed for each condition, and multiple technical replicates within each organoid replicate; shapes on the bar chart correspond to each of the cell lines used (KOLF2 - circles, UCLA6 - square, UCLA1 - triangle). Significance was calculated using the Brown-Forsythe and Welch ANOVA. * indicates p-value < 0.05, ** indicates p-value < 0.01, *** indicates p-value < 0.001. B) snRNA-seq analysis was performed at Week 12 across three organoid lines and all three glucose concentration conditions. UMAPs shown here represent all 81,593 cells passing quality control across lines and conditions. Left: UMAP colored by originating stem cell line (KOLF2-red, UCLA1-green, UCLA6-blue). Middle: UMAP colored by altered glucose conditions (2.5mM-red, 8.75mM-green, 17.5mM-blue). Right: Average cell types across the three stem cell lines. Y-axis represents the fraction of a cell type from the total for a particular glucose condition. Cell types were annotated by mapping to an atlas of human cortical development (Nano et al., 2023). C) UMAP of predicted ID scores from snRNA-seq analysis of glucose manipulation experiment in organoid. This analysis provides a calculation of how well the annotated clusters mapped to the predicted cell type annotation from previous atlas of cortical development. Predicted scores range from 0.2 to 1.0 with a higher score indicating higher likelihood of confidence to predicted cell-type ID. D) UMAPs of key gene neuronal markers for cell types of interest from snRNA-seq analysis. SOX2, HOPX, EOMES, MKI67, DLX6-AS1, GFAP, BCL11B, and SATB2 are neuronal markers for radial glia, oRG, IPC, dividing cells, inhibitory neurons, astrocytes, deep-layer neurons, and upper-layer neurons, respectively. E) Modules of glycolysis, TCA, and PPP genes were examined across glucose conditions highlighting feedback of altered gene expression due to reduced extracellular glucose in organoids. Glycolysis and PPP modules decreased with lower glucose concentration while TCA increased. Module scores were determined by leveraging method from previous meta-atlas study (Nano et al., 2023) to look at gene activity over time. Gene lists were obtained from KEGG pathway analysis. F) Heatmap and bar graph of nucleotide abundance changes in altered glucose levels in organoid shows overall increase of nucleotide intermediates and especially in the level of mono- and diphosphates compared to triphosphates. Metabolite levels in heatmap represent a log_2_ fold change to the baseline/control glucose level (17.5mM). Dark blue or pink values indicate a negative or positive log fold change to 17.5mM glucose, respectively. Gray values represent an absence of data collected for a metabolite from the metabolomics analysis. Black indents at bottom of heatmap separate biological replicates within each timepoint. The y-axis of bar graphs represents the ratio of mono- and di-phosphates over tri-phosphates. Each condition group consisted of three cell lines with three organoid replicates per cell line (KOLF2 - circles, UCLA6 - square, UCLA1 - triangle). Significance was calculated using the Brown-Forsythe and Welch ANOVA. * indicates p-value < 0.05, ** indicates p-value < 0.01, *** indicates p-value < 0.001. G) Bar graph displaying ^13^C-label derived from ^13^C_6_-glucose in AMP, ADP, IMP, and GMP. Each condition group consisted of three cell lines with three organoid replicates per line (KOLF2 - circles, UCLA6 - square, UCLA1 - triangle). Significance was calculated using the Brown-Forsythe and Welch ANOVA. * indicates p-value < 0.05, ** indicates p-value < 0.01, *** indicates p-value < 0.001. H) Bar graph GSH over GSSG ratio indicating oxidative stress among glucose conditions in organoids. Each condition group consisted of three cell lines with three organoid replicates per line (KOLF2 - circles, UCLA6 - square, UCLA1 - triangle). Significance was calculated using the Brown-Forsythe and Welch ANOVA. * indicates p-value < 0.05, ** indicates p-value < 0.01, *** indicates p-value < 0.001.

**Supplemental Figure 7:**
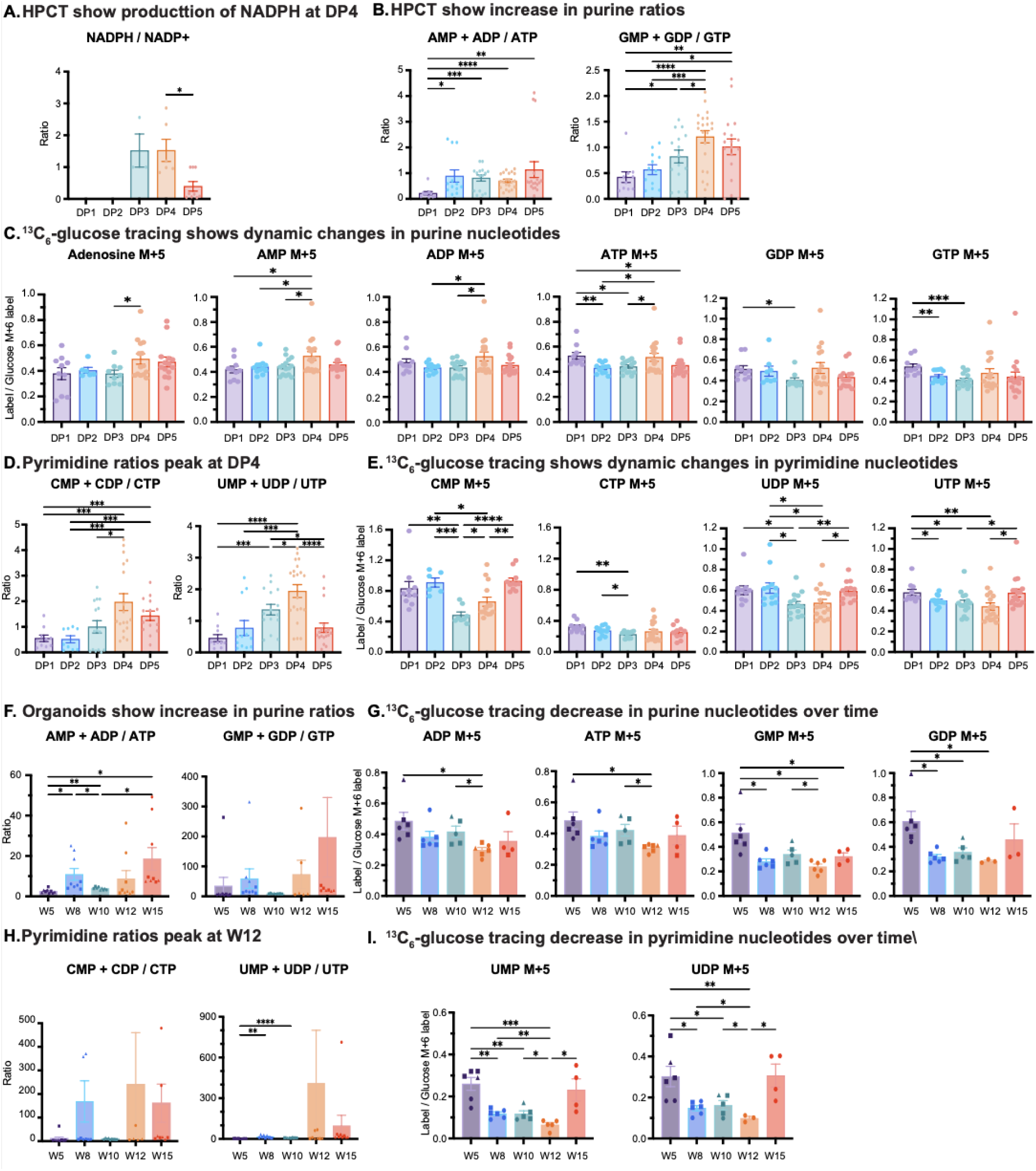
Nucleotide Metabolism Show Dynamic Changes Over the Course of Development. A) A product of PPP activity is the generation of NADPH, we therefore examined the levels of NADPH over NADP+ to determine changes over development in HPCT. Three to five biological replicates of each DP were analyzed, with three to six technical replicates for each sample. Combined metabolite abundance values were calculated by averaging all technical replicates across biological samples within each DP. Dots show technical replicates from each biological sample within DPs. Significance was calculated using the Brown-Forsythe and Welch ANOVA. * indicates p-value < 0.05, ** indicates p-value < 0.01, *** indicates p-value < 0.001. Error bars represent SEM. B) Examination of purine intermediates show increased levels of mono- and di-phosphates over tri-phosphate metabolite abundance shows general increase in HPCT DPs with a peak at DP4. Left graph shows the metabolite abundance ratio of AMP and ADP over ATP, and right graph shows the ratio of GMP and GDP over GDP. C) The amount of ^13^C-label derived from ^13^C_6_-glucose in adenosine, AMP, ADP, ATP, GDP, and GTP shows dynamic changes in purine nucleotides in HPCT across DPs. Glucose tracing graphs display ratio fully labeled isotopologues indicated by M+5. The y-axis measures ratio of ^13^C-label normalized to ^13^C_6_-glucose label (M+6). D) Bar graph shows pyrimidine ratios peak at DP4 in HPCT. Left graph shows the abundance ratio of CMP and CDP over CTP, and the right graph shows the ratio of UMP and UDP over UTP. E) The amount of ^13^C-label derived from ^13^C_6_-glucose in pyrimidines highlights dynamic changes across HPCT DPs. Graphs from left to right display fully labeled isotopologues M+5 of CMP, CTP, UDP, and UTP. Y-axis measures ratio of ^13^C-label normalized to ^13^C_6_-glucose label (M+6). F) Equivalent analysis of changes in nucleotide metabolism in organoid shows general increase in purine ratios over time. Left graph measures the metabolite abundance ratio of AMP and ADP over ATP and the right graph measures the ratio of GMP and GDP over GTP. Three cell lines with three organoid replicates per cell line for each time point were analyzed. Combined metabolite abundance values were calculated by averaging all technical replicates across biological samples within each time point. Dots show technical replicates from each biological sample within each time point (KOLF2 - circles, UCLA6 - square, UCLA1 - triangle). Significance was calculated using the Brown-Forsythe and Welch ANOVA. * indicates p-value < 0.05, ** indicates p-value < 0.01, *** indicates p-value < 0.001. Error bars represent SEM G) The amount of ^13^C-label derived from ^13^C_6_-glucose in ADP, ATP, GMP, and GDP shows general decrease in purine intermediates in organoids across DPs. Graphs measure ratio of fully labeled isotopologues indicated by M+5. The y-axis measures the ratio of ^13^C-label normalized to ^13^C_6_-glucose label (M+6). H) Bar graphs show pyrimidine ratios peak at W12 in organoid metabolic analysis. Left graph shows the abundance ratio of CMP and CDP over CTP, and the right graph shows the ratio of UMP and UDP over UTP. I) ^13^C_6_-glucose tracing displays a general decrease in pyrimidine nucleotides across organoid DPs. Left graph shows the level of fully labeled isotopologues M+5 of UMP and the right graph of UDP. Y-axis measures ratio of ^13^C-label normalized to ^13^C_6_-glucose label (M+6) labeling.

**Supplemental Figure 8:**
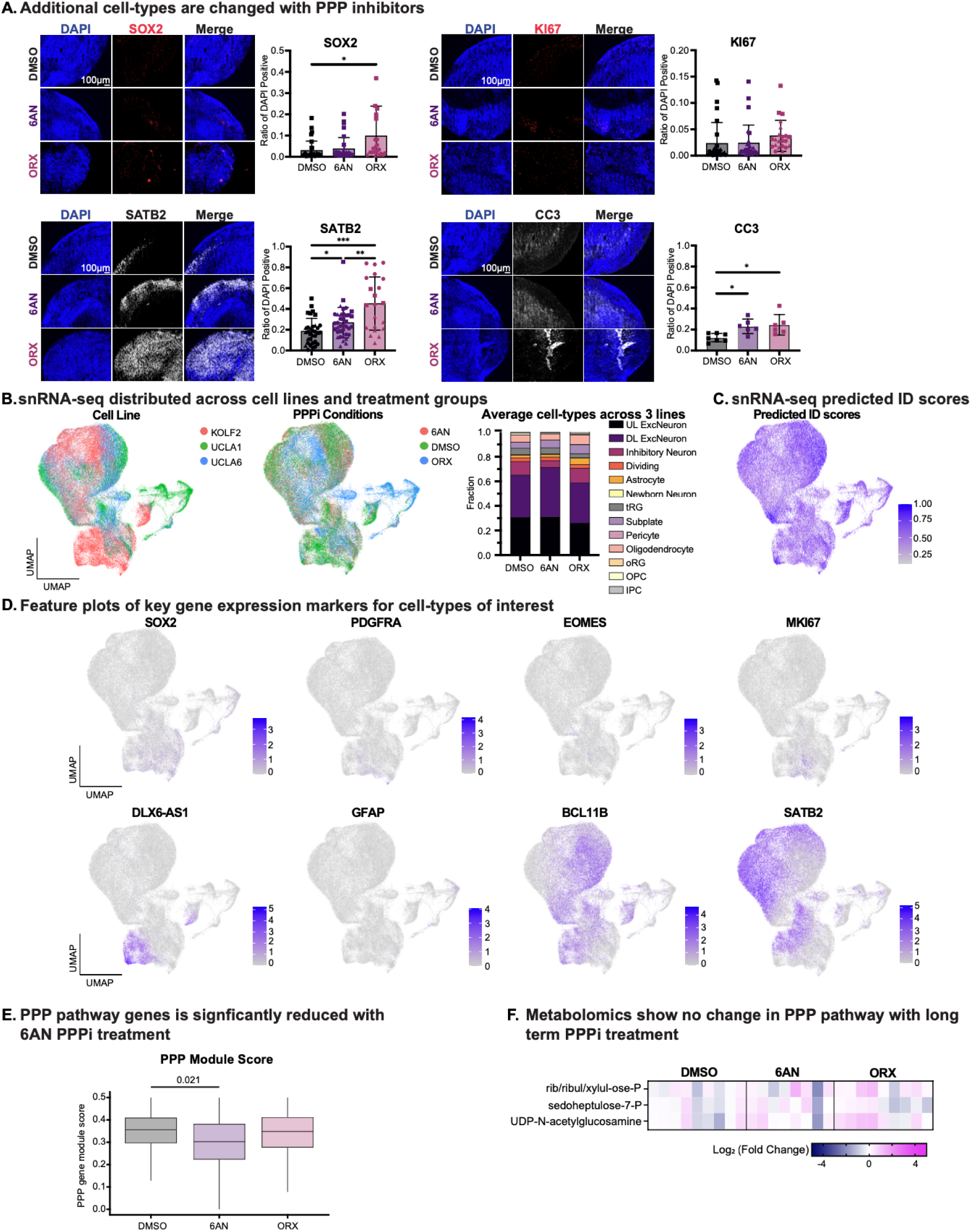
PPP Manipulation Display Cell Type and PPP Gene Signature Changes. A) Additional cell type changes with pharmacological inhibition of PPP measured by IF. A representative image from the KOLF2 line is shown for each of the three condition groups is shown here. Stainings for SOX2, SATB2, KI67, and CC3 mark radial glia, upper-layer neurons, diving cells, and apoptotic cells, respectively. DAPI, shown in blue, marked nuclei. Scale bar is 100 μM and reflects all images. Quantification of the ratio of DAPI positive cells marked by each protein is presented in bar charts to the right, where up to three cell lines with three organoid replicates per cell line were analyzed for each condition; shapes on the bar chart correspond to each of the cell lines used (KOLF2 - circles, UCLA6 - square, UCLA1 - triangle). Significance was calculated using the Brown-Forsythe and Welch ANOVA. * indicates p-value < 0.05, ** indicates p-value < 0.01, *** indicates p-value < 0.001. B) snRNA-seq analysis of W12 organoids across three lines and with pharmacological inhibition of PPP. UMAPs shown here represents all 106,047 cells passing quality control across lines and conditions. Left: UMAP colored by originating stem cell line (KOLF2-red, UCLA1-green, UCLA6-blue). Middle: UMAP colored by PPP inhibitor condition (6AN-red, DMSO-green, ORX-blue). Right: Average cell types across organoids from the three stem cell lines separated by PPP condition. The y-axis represents a fraction of a cell type from the total for a particular glucose condition. Cell types were annotated by mapping to an atlas of human cortical development (Nano et al., 2023). C) UMAP of predicted ID scores from snRNA-seq analysis of PPP manipulation. Predicted scores range from 0.2 to 1.0 with a higher score indicating higher likelihood of confidence to predicted cell type ID. D) UMAPs of key gene expression markers for cell types of interest from snRNA-seq analysis of PPP manipulation in organoids. SOX2, PDGFRA, EOMES, MKI67, DLX6- AS1, GFAP, BCL11B, and SATB2 are neuronal markers for radial glia, oligodendrocyte precursor cells (OPC), IPC, dividing cells, inhibitory neurons, astrocytes, deep-layer neurons, and upper-layer neurons, respectively. E) Modules activity of PPP genes was observed in PPP inhibitor conditions and show feedback of reduced PPP gene expression changes with 6AN inhibitor. Module scores were determined by leveraging method from previous meta-atlas study (Nano et al., 2023) to look at gene activity over time. Gene lists were obtained from KEGG pathway analysis. Significance was determined by performing a t-test. F) Heatmap of metabolomics analysis of PPP intermediates shows no change with long term PPP inhibitor treatment at W12 harvest. Metabolite levels in the heatmap represent a log2 fold change to DMSO control. Analysis includes three cell lines with three organoids per cell line.

**Supplemental Figure 9:**
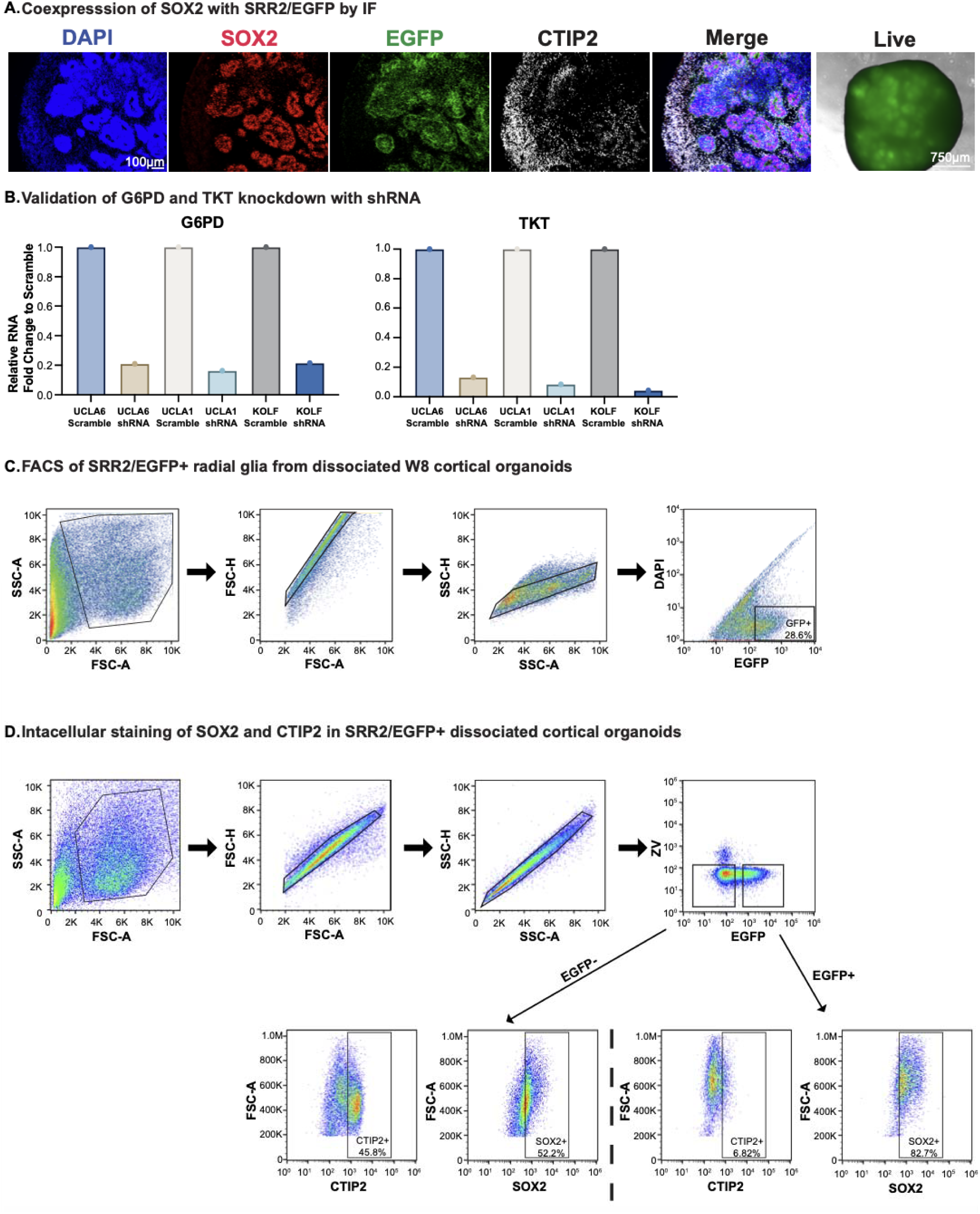
Workflow to Isolate Radial Glia and Introduce shRNAs Targeting PPP Enzymes. A) SRR2 enhancer appropriately enriches for selection of radial glia from organoid as validated with immunofluorescent images that show high overlap of SOX2 and SRR2/EGFP with minimal overlap with CTIP2. A representative image from the KOLF2 line is shown here for SOX2, CTIP2, EGFP, and live. Staining for SOX2 (red) was used as a proxy for radial glia progenitor identity, whereas staining for CTIP2 (white) labeled emerging deep layer neuron populations. DAPI, shown in blue, marked nuclei. Scale bar is 100 μM and reflects all images. Right-most image shows live SRR2/EGFP imaging in cortical organoids and displays strong signal in rosettes where radial glia is most present. B) Validation of PPP genes knockdown with appropriate shRNA by RT-qPCR shows strong decreases in RNA relative to scramble control across organoids from three stem cell lines. Organoids were dissociated and plated down on coated plated and infected with lentivirus for shRNA production targeting either G6PD or TKT before collection of RNA 72 hours post infection. Gene expression quantification is displayed as relative RNA fold change to scramble for G6PD (left) and TKT (right). C) FACS of SRR2/EGFP+ radial glia from dissociated W8 cortical organoids for subsequent plating onto coated plates and knockdown of PPP genes. Several organoids across three lines were dissociated and prepped to achieve appropriate cell density for subsequent plating. FACS show selection of cells and debris removal based on forward and side scatter area, followed by two-step doublet removal using forward and side scatter height and area. Live cells were selected based on negative signal from DAPI and SRR2+ radial glia cells were selected based on log fold signal intensity of EGFP. D) Further validation of SOX2 enrichment in SRR2/EGFP+ cells by intracellular staining of SOX2 and CTIP2 and subsequent flow cytometry of dissociated organoids. TOP: density plots show selection of cells and debris removal based on forward and side scatter area, followed by two-step doublet removal using forward and side scatter height and area. Live cells were selected based on negative signal from zombie-violet, an extracellular marker of cell death, and SRR2+ radial glia cells were selected based on log fold signal intensity of EGFP. BOTTOM: left two density plots display positive signal of CTIP2 and SOX2 from the EGFP- gated organoid cells and right two density plots show increased SOX2 signal and lower CTIP2 signal from EGFP+ gate.

**Supplemental Figure 10:**
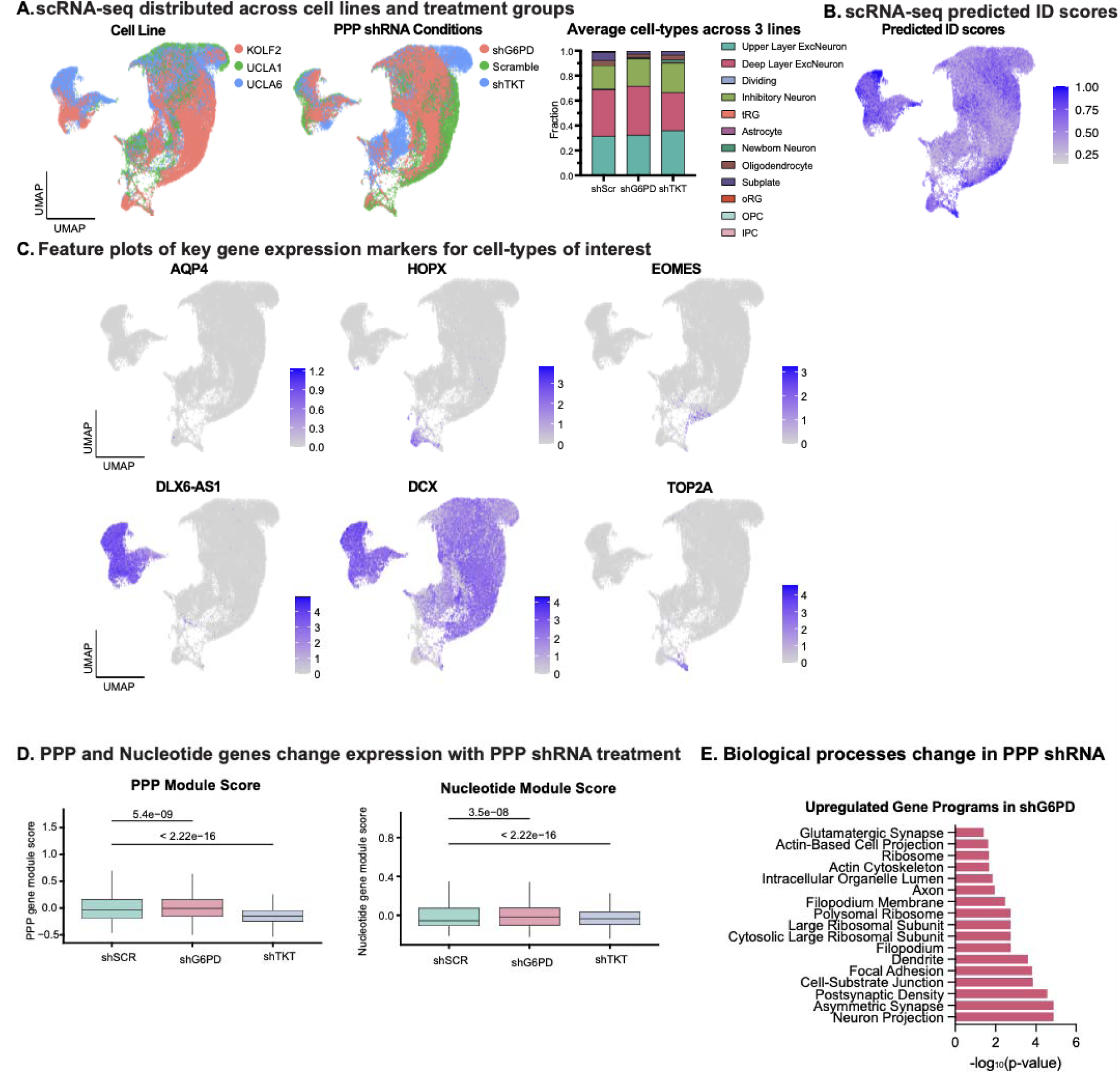
Genetic Knockdown of PPP Enzymes Impact Cell Types and Metabolic Signatures. A) scRNA-seq analysis from genetic knockdown of PPP genes using shRNAs targeting G6PD or TKT in organoids. UMAPs reflect all 51,105 cells passing quality control across lines and conditions. Left: UMAP colored by originating stem cell line (KOLF2-red, UCLA1-green, UCLA6-blue). Middle: UMAP colored by PPP knockdown conditions (shG6PD-red, Scramble-green, shTKT-blue). Right: Average cell types across organoid cells from three stem cell lines separated by PPP knockdown condition. The y-axis represents a fraction of a cell type from the total for a particular glucose condition. B) UMAP of predicted ID scores from scRNA-seq analysis of genetic PPP manipulation. Predicted scores range from 0.2 to 1.0 with a higher score indicating higher likelihood of confidence to predicted cell-type ID. C) UMAPs of key gene expression markers for cell types of interest from snRNA-seq analysis of PPP enzyme knockdown in organoid. AQP4, HOPX, EOMES, DLX6-AS1, DCX, TOP2A are markers for astrocyte, oRG, IPC, inhibitory neurons, newborn neurons, and dividing cells, respectively. D) Modules activity of PPP genes in knockdown conditions display dynamic gene expression feedback responses to changes in PPP enzyme activity. Module scores were determined by leveraging a method from a previous meta-atlas study (Nano et al., 2023) to look at gene activity over time. Gene lists were obtained from KEGG pathway analysis. Significance was determined by t-test analysis. Gene Ontology Biological Process analysis of the genes upregulated in the shG6PD condition highlight gene programs related to neuronal development.

